# Dynamic measurement of cytosolic pH and [NO_3_^-^] uncovers the role of the vacuolar transporter AtCLCa in the control of cytosolic pH

**DOI:** 10.1101/716050

**Authors:** Elsa Demes, Laetitia Besse, Béatrice Satiat-Jeunemaitre, Sébastien Thomine, Alexis De Angeli

**Affiliations:** Institute for Integrative Biology of the Cell (I2BC), CEA, CNRS, Univ. Paris-Sud, Université Paris-Saclay, 91198, Gif-sur-Yvette cedex, France

**Keywords:** Arabidopsis, vacuole, biosensor, nitrate, exchanger, chloride

## Abstract

Ion transporters are key players of cellular processes. The mechanistic properties of ion transporters have been well elucidated by biophysical methods. Meanwhile the understanding of their exact functions in the whole cell homeostasis is limited by the difficulty to monitor their activity *in vivo*. The development of biosensors to track subtle changes in intracellular parameters provides an invaluable key to tackle this challenging issue. Here, we adapted the use of a dual biosensor using guard cells as experimental model to visualize the impact on the cytosol of anion transport from intracellular compartments. To image the activity of AtCLCa, a vacuolar NO_3_^-^/H^+^ exchanger regulating stomata aperture in *Arabidopsis thaliana*, we expressed a genetically encoded biosensor, ClopHensor allowing monitoring the dynamics of cytosolic anion concentration and pH. We first show that ClopHensor is not only a Cl^-^ but also a NO_3_^-^ sensor. We were then able to unravel and quantify the variations of NO_3_^-^ and pH in the cytosol. Our data show that AtCLCa activity modifies cytosolic pH and NO_3_^-^, demonstrating that the transport activity of a vacuolar exchanger has a profound impact on cytosolic homeostasis. We propose that a major function of this endomembrane transporter is to adjust cytosolic conditions to cellular needs. This opens a novel perspective on the function of intracellular transporters of the CLC family in eukaryotes: not only controlling the intra organelle lumen but also actively modifying cytosolic conditions.

**Significance:** Intracellular transporters are key actors in cell biological processes. Their disruption causes major physiological defects. The role of intracellular ion transporters is usually seen through an “intra organelle” lens, meanwhile their potential action on cytosolic ion homeostasis is still a black box. The case of a plant CLC is used as a model to uncover the missing link between the regulation of conditions inside the vacuole and inside the cytosol. The development of an original live imaging workflow to simultaneously measure pH and anion dynamics in the cytosol reveals the role of an *Arabidopsis thaliana* CLC, AtCLCa, in the modification of cytosolic pH. Our data highlight an unsuspected function of endomembrane transporters in the regulation of cytosolic pH.

## Introduction

The fluxes of ions between cell compartments are ruled by membrane proteins forming ion channels, exchangers, symporters and pumps. Defects in the transport systems residing in intracellular membranes results in major physiological failures at the cellular and the whole organism level. The localization of transport systems in intracellular membranes prevents the use of *in vivo* electrophysiological approaches, considerably limiting our understanding of their subcellular functions. Among the different families of ion transporters identified, the CLC (Chloride Channel) family has been widely investigated in the last decades (1). CLCs form a family of membrane proteins present in all organisms (1). The members of the CLC family function as anion channels or anion/H^+^ exchangers sharing a similar structural fold (2, 3). In eukaryotes, all the CLCs localized in intracellular membranes behave as anion/H^+^ exchangers. In mammals, mutations in intracellular CLCs lead to severe genetic diseases affecting bones, kidneys and brain (1). In plants, CLCs regulate nutrient storage, photosynthesis and participate to drought and salt stress tolerance (4–10). In the last decades a wide number of studies addressed the biophysical properties of intracellular CLCs and provided a solid ground to understand the transport mechanisms of these exchangers (11–15). However, we still lack a molecular interpretation of the role of the CLC exchangers within the cells, preventing a full understanding of the defects caused by their mutations (1).

Plant guard cells (GCs) constitute an appropriate experimental model to unravel CLC functions at subcellular level. In plants, the guard cells (GCs) are specialized cells gating the stomata pores at the leaf surface. Their biological function relies on the regulation of the ion transport systems residing in the plasma membrane (PM) and vacuolar membrane (VM) (16–18). The VM delimits the largest intracellular compartment of GCs, the vacuole (16, 19). Stomata regulate gas exchanges between the photosynthetic tissues and the atmosphere, simultaneously controlling water loss by evaporation. Two GCs delimit and regulate the stomata pore aperture according to the environmental conditions. The regulation of the stomata pore aperture is based on the capacity of GCs to change their turgor pressure and, consequently, their shape. Increase and decrease of the turgor pressure in GCs open and close the stomata, respectively. Turgor changes in GCs depends on the accumulation/release of ions into/from the vacuole. Therefore, vacuolar ion transporters are key actors of stomata responses. Over the last decades, the identification of a growing number of genes encoding ion transporters and channels in the VM of GCs highlighted the importance of the intracellular transport systems selective for NO_3_^-^, Cl^-^ and malate^2-^ (6, 7, 20–23). Anion channel and transporter families such as SLAC/SLAH (Slow Activating Anion Channels), ALMT (Aluminum Activated Malate Transporter) and the Chloride channel (CLC) strongly influence GCs function and stomata responses to environmental changes (6, 20–22, 24–27). However, the observed GCs phenotypes and the biophysical characteristics of these ion transport systems can be somehow difficult to reconcile (6, 7, 19, 20, 25, 28). The vacuolar CLC AtCLCa is illustrative of this difficulty. AtCLCa is known to act as a 2 NO_3_^-^/1H^+^ exchanger driving the accumulation of NO_3_^-^ into the vacuole (5, 29), suggesting a role in stomata opening. However, analysis of GCs responses from plants knocked-out in *AtCLCa* gene revealed that the AtCLCa exchanger is not only involved in light-induced stomata opening but also in ABA-induced stomata closure (6). This intriguing dual role questions the molecular interpretation of the subcellular role of AtCLCa.

Being anion/H^+^ exchangers, intracellular CLCs are expected to induce simultaneous modifications of [NO_3_^-^, Cl^-^] and pH in both the lumen of intracellular compartments and the cytosol. However, so far, only their role in regulating luminal-side parameters has been investigated both in plants (using isolated plant vacuoles (30)), and in mammalian (in lysosomes and endosomes (13, 15, 31)). In mammals, CLC-5 was shown to participate to acidification of endosomes (31), while CLC-7 activity was associated only to modest and controversial changes of lysosomal pH (15, 31). In both cases, the link between luminal acidification and the severe phenotypes in the corresponding knock-out mice was not established (15, 31). In plants, no report of a role of a CLC in vacuolar pH regulation was so far demonstrated *in vivo*. Here, we hypothesized that AtCLCa activity affects cytosolic parameters in addition to its well-documented role in anion accumulation inside vacuoles. We therefore aimed to visualize whether the activity of an intracellular CLC like AtCLCa induces changes in the cytosolic pH and [NO_3_^-^, Cl^-^] dynamics in living GCs.

In order to be able to detect simultaneously the subtle changes in cytosolic pH and anion concentration induced by the activity of an intracellular transporter, we introduced the genetically encoded biosensor ClopHensor in GCs as an experimental model. ClopHensor is a ratiometric biosensor originally developed in mammalian cells with spectroscopic properties allowing to measure [Cl^-^] and pH in parallel (32). Our results demonstrated that ClopHensor allows simultaneous measurement of the cytosolic pH and [Cl^-^] (32) and, additionally, [NO_3_^-^] in GCs. NO_3_^-^ is a highly relevant anion in plant cells, defining ClopHensor as the first biosensor for NO_3_^-^. Arabidopsis GCs expressing ClopHensor in the cytosolic compartment were used to visualize *in vivo* the subcellular effects of the activity of the NO_3_^-^/H^+^ exchanger AtCLCa. We monitored by live Confocal Laser Scanning Microscope (CLSM) the evolution of [Cl^-^]_cyt_ or [NO_3_^-^]_cyt_ in parallel with pH_cyt_. We developed a specific image analysis workflow to measure the fluorescence ratios of interest in GCs. A comparative study between wild-type and mutant GCs knocked-out for AtCLCa shows that the vacuolar exchanger AtCLCa controls not only the kinetics of cytosolic [NO_3_^-^] changes, but also actively participates in the control of cytosolic pH. These results highlight an unexpected role of CLCa in the regulation of cytosolic pH. Further, they open a new perspective on the cellular functions of intracellular transporters in GCs that may provide an integrated framework to understand the function of intracellular CLCs in other eukaryotic cells.

## Results

### In vitro assays reveal a strong affinity of ClopHensor for NO_3_^-^

In contrast with mammalian cells, several anionic species are present in the millimolar range in plant cells (4, 33). Therefore, we investigated the sensitivity of ClopHensor to Cl^-^, NO_3_^-^, PO_4_^3-^, malate^2-^ and citrate^3-^, the main anions present in the model plant *Arabidopsis* (4). We used recombinant ClopHensor proteins bound to sepharose beads (Fig. 1A), and recorded the fluorescence upon exposure to a range of anions by CLSM after excitation at 561, 488 and 458 nm. The imaging ratio R_Anion_ (F_458_/F_561_) was calculated from the ratio of the fluorescence intensity images after excitation at 458 nm (F_458_) and 561 (F_561_) to estimate the effect of anions on ClopHensor (see Supplementary Material for calculation R_Anion_). No significant difference in R_Anion_ was observed between control and 30 mM PO_4_^3-^, malate^2-^ and citrate^3^ (Fig. 1A). Meanwhile, we found that ClopHensor was sensitive to Cl^-^ and, remarkably, also to NO_3_^-^ (Fig.1A). ClopHensor displayed a higher affinity to NO_3_^-^ (K_d_^NO3^=5.3±0.8 mM at pH=7) than to Cl^-^ (K_d_^Cl^ =17.5±0.5 at pH=6.8) (Fig. 1B). The usefulness range of ClopHensor for Cl^-^ was between 2.5 and 125 mM (at pH 6.8) and for NO_3_ ^-^ between 0.6 and 32 mM (at pH 7) (Fig. 1B). To test the pH sensitivity of ClopHensor in our *in vitro* assay, we calculated the imaging ratio R_pH_ (F_488_/F_458_) (see Supplementary Material for calculation R_pH_). In agreement with previous reports (32), we found a strong response of RpH to pH variations with a steep dynamic range of 9 fold change within 2 pH units, and a pK_a_= 6.98±0.09, (Fig. 1D). The usefulness range of ClopHensor was estimated for pHs between 6.1 and 7.9 *in vitro* (Fig. 1D). Neither the binding of NO_3_^-^ nor that of Cl^-^ did modify significantly the pH sensitivity of ClopHensor (Fig. S1C), confirming its robustness as an anion biosensor to be used in plant cells.

**Fig. 1.**
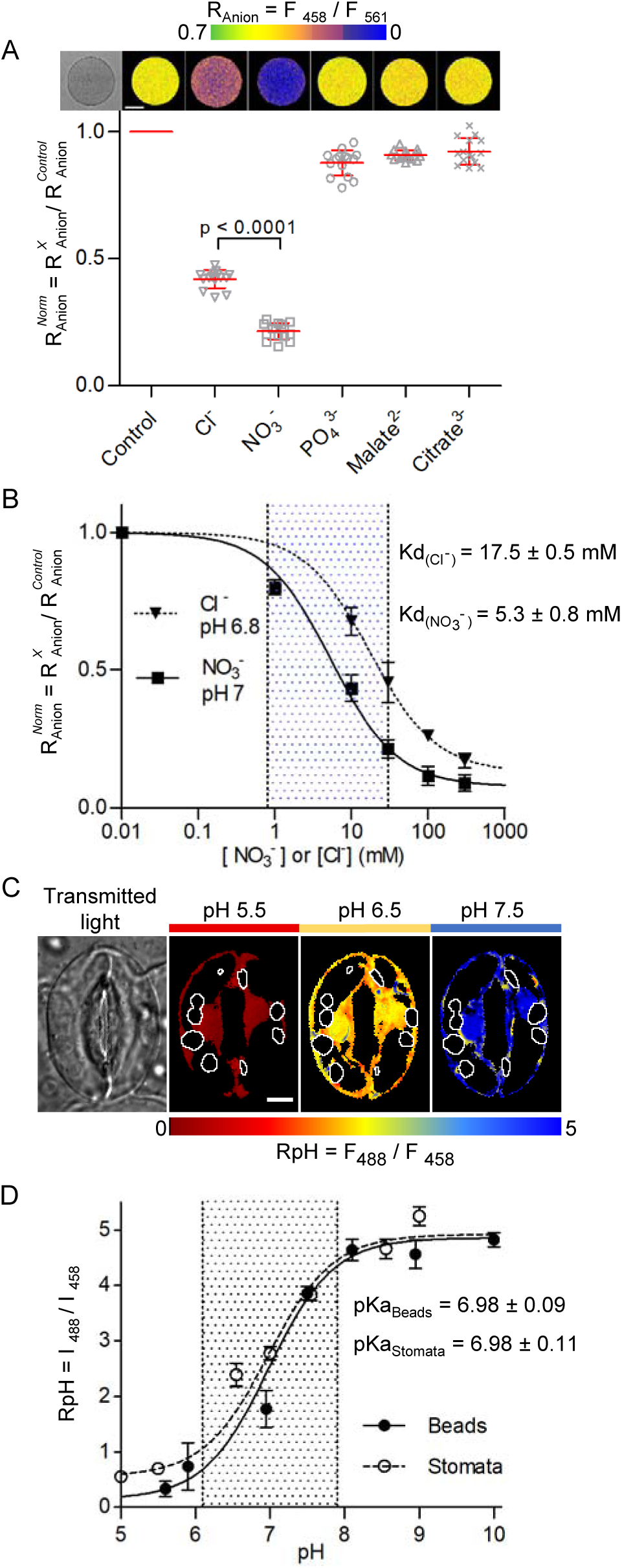
ClopHensor is sensitive to NO_3_^-^, Cl^-^ and pH. (A) *In vitro* ratio imaging of sepharose beads decorated with ClopHensor. *Upper panel*, false colour images of representative beads in presence of 30 mM Cl^-^, NO_3_^-^, PO_4_^3-^, malate^2-^ and citrate^3-^ displaying the fluorescence ratio R_Anion_ (R_Anion_ = F_458_/F_561_; scale bar 50 µm). *Lower panel*, R_Anion_ normalized to R_Anion_ in control conditions in presence of 30 mM Cl^-^, NO_3_^-^, PO_4_^3-^, malate^2-^ and citrate^3-^ (mean value ± SD, n≥15 beads for each condition) The bracket indicates statistically significant difference (T-test). (B), *in vitro* dose response analysis of R_Anion_ in presence of Cl^-^ or NO_3_^-^ from 0 to 300 mM at pH 6.8 and 7, respectively (mean value ± SD, n=15 beads for each condition). The dissociation constant Kd was obtained fitting the data with equation S2. Dotted area, usefulness range for NO_3_^-^ (0.6 to 30 mM). (C) *In vivo* ratio imaging of Arabidopsis stomata expressing ClopHensor. False colour images of a representative stomata showing the fluorescence ratio RpH (RpH= I_488_/I_458_) upon sequential exposure to extracellular NH_4_-acetate buffers at pH 5.5, 6.5 and 7.5. Left to right transmitted light, false colour images of RpH at pH 5.5, 6.5 and 7.5. Top horizontal bar indicates the pH of the buffer based on the false colour scale (lower horizontal bar). White contours represent the chloroplast position. Scale bar = 5 µm. (D) plot of RpH vs pH shows that the titration curves of ClopHensor *in vivo* (stomata, black circles; n≥10) and *in vitro* (sepharose beads, empty circles; n≥15) are comparable. Data were fitted with equation S1, the dotted area shows the useful pH range (6.1 to 7.9) of ClopHensor. All error bars represent the standard deviation.

### ClopHensor is a robust and sensitive sensor of cytosolic pH in *Arabidopsis thaliana* guard cells

We generated transgenic Arabidopsis plants (ecotype Col-0) expressing ClopHensor in the cytosol and the nucleoplasmic compartments under the control of the Ubiquitin10 promoter (*pUB10*:*ClopHensor)*. The expression of ClopHensor did not affect the development of the plants, indicating that its expression did not significantly interfere with the amount of anions available in the cytosol for cellular metabolism (Fig. S2). To measure the pH sensitivity of ClopHensor in living GCs, stomata from *pUBI10*:*ClopHensor* were sequentially exposed to NH_4_-acetate based buffers to clamp the pH_cyt_ at defined values between 5 and 9 (Fig. 1C, D). We found that ClopHensor sensitivity to pH *in vivo* and *in vitro* are very similar. The mean <RpH_cyt_>, calculated from each pixel in the stomata (Fig. 1D), showed that the pH titration curve of ClopHensor in GCs mirrored the *in vitro* assay (Fig. 1D). The *in vivo* pKa (6.98±0.11) and the dynamic range of ClopHensor match the values measured *in vitro* (Fig. 1D). These findings demonstrate that: i) ClopHensor is a reliable reporter for intracellular pH changes in GCs; ii) the cytosolic environment does not affect ClopHensor properties; iii) ClopHensor sensitivity range is appropriate to measure cytosolic pH in GCs.

### Settings and design of the experimental workflow in GCs

The data we obtained open to the possibility to measure the dynamic evolution of [NO_3_^-^]_cyt_, [Cl^-^]_cyt_ and pH_cyt_ *in vivo*. This provides a unique opportunity to disclose in living cells how ion fluxes across the PM and the VM of GCs affect cytosolic conditions. In order to precisely quantify [NO_3_^-^]_cyt_, [Cl^-^]_cyt_ and pH_cyt_ in GCs we optimized the fluorescence acquisition protocol in GCs expressing ClopHensor (Fig. S3 and S6), and determined the temporal window to set up our experiments. First, to maximize the collected fluorescence and minimize photo-damage by the laser, we selected stable transgenic lines expressing *pUBI10*:*ClopHensor* with high fluorescence in GCs after excitation at 561, 488 and 458 nm. Second, to precisely quantify [NO_3_^-^]_cyt_, [Cl^-^]_cyt_ and pH_cyt_, we excluded the fluorescent signals emitted by chloroplasts. Therefore, we developed an image processing workflow to accurately measure ClopHensor fluorescence in the cytosol of plant cells (Fig. S6).

To derive the [NO_3_^-^]_cyt_, [Cl^-^]_cyt_ and pH_cyt_ in GCs, we used the calculation described in Arioso et al. (2010) (Supplementary Information Text). To obtain a quantitative estimation of the changes in [NO_3_^-^]_cyt_ and [Cl^-^]_cyt_ induced by the applied treatments, we determined *in vivo* the R_Anion_ ratio in the absence of NO_3_^-^ and Cl^-^ (i.e. R^0^). R^0^ is required to calculate the actual concentration of Cl^-^ and NO_3_^-^ in the cytosol (Supplementary Information Text). To this aim, we set up experimental conditions where the initial endogenous [NO_3_^-^]_cyt_ and [Cl^-^]_cyt_ were below the sensitivity threshold of ClopHensor: we grew *in vitro pUB10*:*ClopHensor* plants in a NO_3_^-^-free medium (0 NO_3_^-^ medium) and determined the whole plant [NO_3_^-^] and [Cl^-^] at different days after germination (DAG) (Table S1). We found that the endogenous content of NO_3_^-^ and Cl^-^ was decreasing after germination. At 14 DAG, Cl^-^ was no longer detectable, meanwhile [NO_3_^-^] was below the sensitivity threshold of ClopHensor (i.e. 0.6 mM). Subsequently, based on these data, we imaged stomata from *pUB10*:*ClopHensor* plants grown for 14 days on a NO_3_^-^-free medium and measured a mean ratio R^0^_Anion_ of 0.56±0.07 (n=29 stomata) (Fig. S3).

### Dynamic measurements of cytosolic NO_3_^-^, Cl^-^ and pH in Arabidopsis GCs

We then challenged ClopHensor for the simultaneous detection in GCs of [NO_3_^-^]_cyt_, [Cl^-^]_cyt_ and pH_cyt_ changes upon extracellular NO_3_^-^ or Cl^-^ supply/removal (Fig. 2). The experimental design was based on the application of different extracellular conditions in a sequence of 5 steps (Fig. 2). GCs were, (1) perfused with NO_3_^-^-free medium to establish the ratio R^0^ for each stomata ; (2) exposed to 30 mM KNO_3_ to observe [NO_3_^-^]_cyt_ changes ; (3) washed-out with NO_3_^-^-free medium ; (4) exposed to 30 mM KCl to observe [Cl^-^]_cyt_ changes ; (5) washed-out again with NO_3_^-^-free medium. We applied 30 mM KNO_3_ or KCl as these concentrations are commonly used in stomata aperture assays (7, 34). To perform a full experiment, we imaged GCs for 190 min, each stomata was imaged every 4 minutes with a sequential excitation at 561, 488 and 458 nm. ClopHensor was not significantly affected by photo-bleaching over the whole duration of the experiments, as the fluorescence intensity recorded in NO_3_^-^-free media was not altered after 190 minutes of illumination (Fig. S4). Raw data suggested striking variations of the mean fluorescence intensity recorded after excitation at 488 nm and 458 nm when NO_3_ ^-^ was added to –or washed-out from–the extracellular medium, meanwhile Cl^-^ addition had less pronounced effects (Fig. S4). Ratiometric images for R_pH_ and R_Anion_ were established from the fluorescence intensity images (respectively Fig. 2A and 2C). The ratiometric maps for R_pH_ and R_Anion_ were then used to compute the mean pH_cyt_, (Fig. 2B), and the mean [NO_3_^-^]_cyt_ and [Cl^-^]_cyt_ for each stomata(Fig. 2D, E). The results confirmed that, differently from NO_3_^-^, no significant difference was detected after Cl^-^ addition, as [Cl^-^]_cyt_ stood below the sensitivity threshold of ClopHensor (Fig. 2E).

**Fig. 2.**
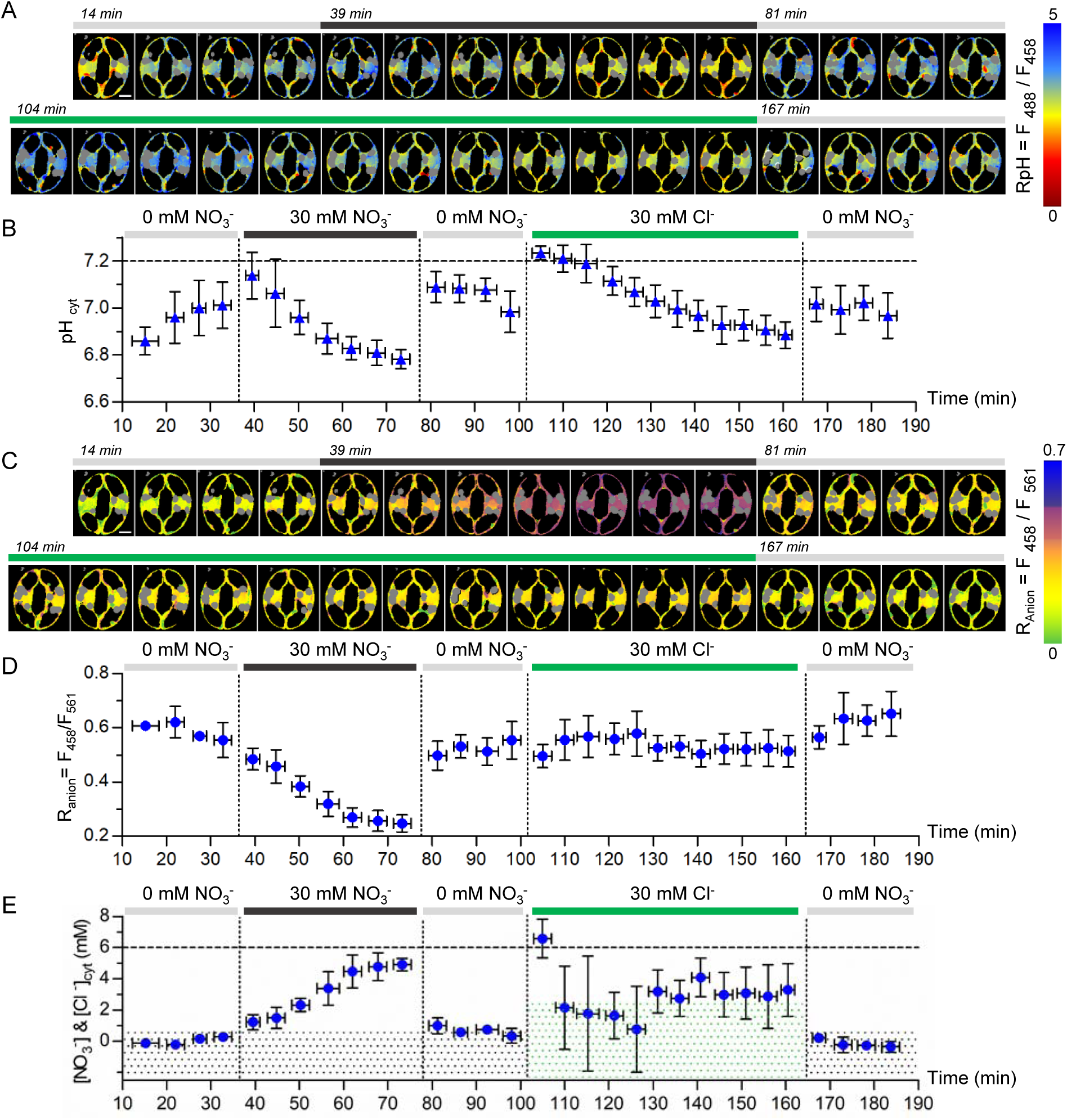
ClopHensor reveals the dynamics of cytosolic pH, NO_3_^-^ and Cl^-^ in Arabidopsis stomata. Representative false colour ratio images of R_pH_ (A) and R_Anion_ (C), at different time points of a stomata sequentially exposed to different extracellular conditions: NO_3_^-^ free medium (0 mM NO_3_^-^, light grey horizontal bar), 30 mM KNO_3_ (dark grey horizontal bar) and 30 mM KCl (green horizontal bar). Scale bar = 5 µm. (B) cytosolic pH was quantified at each time point from the corresponding RpH images. quantification of the temporal evolution of the R_Anion_ in the cytosol of GCs. (E) cytosolic [NO_3_^-^] and [Cl^-^] were calculated at each time point. (B and E), cytosolic pH and [NO_3_^-^] change simultaneously upon extracellular exposure and removal of 30mM KNO_3_. The dotted zone indicates the sensitivity threshold of ClopHensor for NO_3_^-^ (black) and Cl^-^ (green). (B, C and D), data are mean value ± standard deviation of 3 independent stomata. Supplementary Information shows the workflow for the calculation of cytosolic pH and [NO_3_^-^] or [Cl^-^] in (B) and (E). Horizontal error bars represent the time interval of 4 minutes for the sequential imaging of stomata. Vertical dotted lines indicate changes of extracellular conditions. The horizontal dashed lines serves as references for pH 7.2 and [anion]_cyt_ = 6 mM for comparison with figures 3 and 4. Grey areas in (A) and (B) show the position of the chloroplasts and were excluded from the analysis.

In addition, the comparison of pH_cyt_ (Fig. 2B) and [NO_3_^-^]_cyt_ changes (Fig. 2D) over the experiment time suggests a link between NO_3_^-^ transport and pH modification. Initially (step1), in the NO_3_^-^-free medium, the pH_cyt_ was 6.86±0.06. Within 35 minutes it increased and stabilized to 7.01±0.10, while [NO_3_^-^]_cyt_ was below the limit of detection of ClopHensor (i.e. 0.6 mM) (Fig. 2E). Upon addition of 30 mM extracellular KNO_3_ (step 2), the [NO_3_^-^]_cyt_ increased to 4.91±0.40 mM, as expected. In parallel, the pH_cyt_ decreased to 6.78±0.04; both pH_cyt_ and [NO_3_^-^]_cyt_ reached a plateau within 20-30 minutes, suggesting a coordination between the 2 parameters. At step 3, unexpectedly, upon removal of KNO_3_, both pH_cyt_ and [NO_3_^-^]_cyt_ dropped back to the initial values in less than 4 minutes. Finally (step 4), when the stomata were exposed to 30 mM KCl, a progressive decrease of pH_cyt_ was recorded but with a kinetic slower than with 30 mM KNO_3_ (Fig. 2B).

As a whole, these data demonstrate that ClopHensor enables to simultaneously monitor *in vivo* the variations in [Cl^-^]_cyt_ or [NO_3_^-^]_cyt_ and pH_cyt_ at a subcellular resolution. In the tested conditions [Cl^-^]_cyt_ increased to values close to the limit of detection of ClopHensor for Cl^-^ (i.e. 2.5 mM) (Fig. 2E), therefore implying that, in our experimental setting ClopHensor was measuring essentially cytosolic NO_3_^-^ variations. Notably, cytosolic NO_3_^-^ and pH changes appear to be concerted suggesting that they are generated by a common mechanism.

### A vacuolar ion transporter controls cytosolic anion and pH homeostasis

The finding that ClopHensor can measure the dynamic changes of [NO_3_^-^]_cyt_ and pH_cyt_ in GCs opens the possibility to visualize the activity of intracellular ion transport systems in living cells. To test this, we addressed the role of the vacuolar 2NO_3_^-^/1H^+^ exchanger AtCLCa in regulating cytosolic NO_3_^-^ and pH homeostasis. AtCLCa is known to allow the uptake of NO_3_^-^ into the vacuole driven by H^+^ extrusion into the cytosol (5, 35). Therefore, based on its biophysical properties, AtCLCa may be involved in the [NO_3_^-^]_cyt_ and pH_cyt_ responses we measured in Fig. 2.

To assess this possibility, we generated *clca-3* knockout mutant expressing ClopHensor by crossing *clca-3* with a wild type *pUBI10:ClopHensor* line. Patch-clamp experiments performed on vacuoles isolated from wild type and *clca-3 pUBI10:ClopHensor* confirmed that *clca-3* plants expressing *pUBI10:ClopHensor* were defective in vacuolar NO_3_^-^ transport activity (Fig. S7). We then compared the dynamic changes of [NO_3_^-^]_cyt_ and pH_cyt_ in stomata from wild type and *clca-3 pUBI10:ClopHensor* plants (Fig. 3 and 4). Since AtCLCa is highly selective for NO_3_^-^ over Cl^-^ we performed experiments applying extracellular KNO_3_ only. Again, we designed experiments divided in 5 steps. GCs from wild type and *clca-3* were, (1) perfused with NO_3_^-^-free medium to establish the ratio R^0^ of each stomata; (2) perfused with 10 mM KNO_3_; (3) washed-out with NO_3_^-^-free medium ; (4) perfused with 30 mM KNO_3_; (5) washed-out with NO_3_^-^-free medium.

**Fig. 3.**
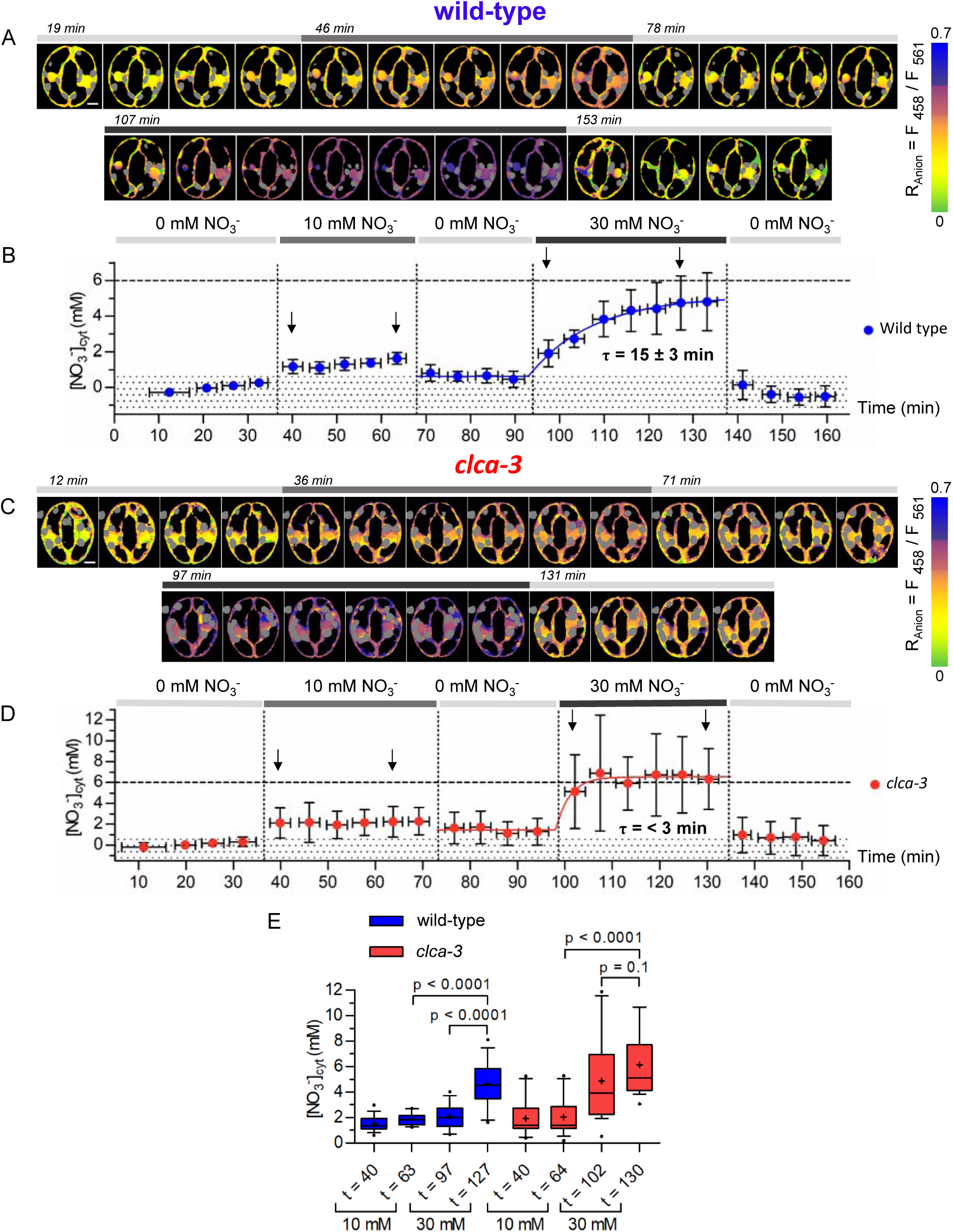
The vacuolar NO_3_^-^/H^+^ exchanger AtCLCa controls [NO_3_^-^]_cyt_ in Arabidopsis stomata. Representative false colour ratio images of R_Anion_ from (A) wild-type and (C) *clca-3* stomata at different time points. Stomata were sequentially exposed to different extracellular conditions as indicated (top horizontal bar). Scale bar = 5 µm. Mean [NO_3_^-^]_cyt_ at each time point in (B) wild type (n=8) and (D) *clca-3* stomata (n=15). The [NO_3_^-^]_cyt_ depends on the applied extracellular [KNO_3_] (top horizontal bar). Note the progressive increase of [NO_3_^-^]_cyt_ after application of 30mM KNO_3_ in wild type (B) and the immediate increase of [NO_3_^-^]_cyt_ in *clca-3* (D). The horizontal bars show the time interval in which stomata were imaged. Vertical error bars are standard deviation. Dotted areas illustrate the sensitivity threshold for NO_3_^-^ of ClopHensor. Vertical dotted lines indicate changes of extracellular conditions. The horizontal dashed lines serves as reference for [anion]_cyt_ = 6 mM for comparison between Figure 2E, 3B and 3D. Grey areas in (A) and (B) show the position of the chloroplasts and were excluded from the analysis. (E) Box plots of the [NO_3_^-^]_cyt_ at different time points (black arrows in B and C). The brackets indicate statistically significant differences. Blue boxes: wild type (n=17); red boxes: *clca-3* (n=15) stomata. Whiskers show the 10-90% percentile; the cross indicates the mean.

**Fig. 4.**
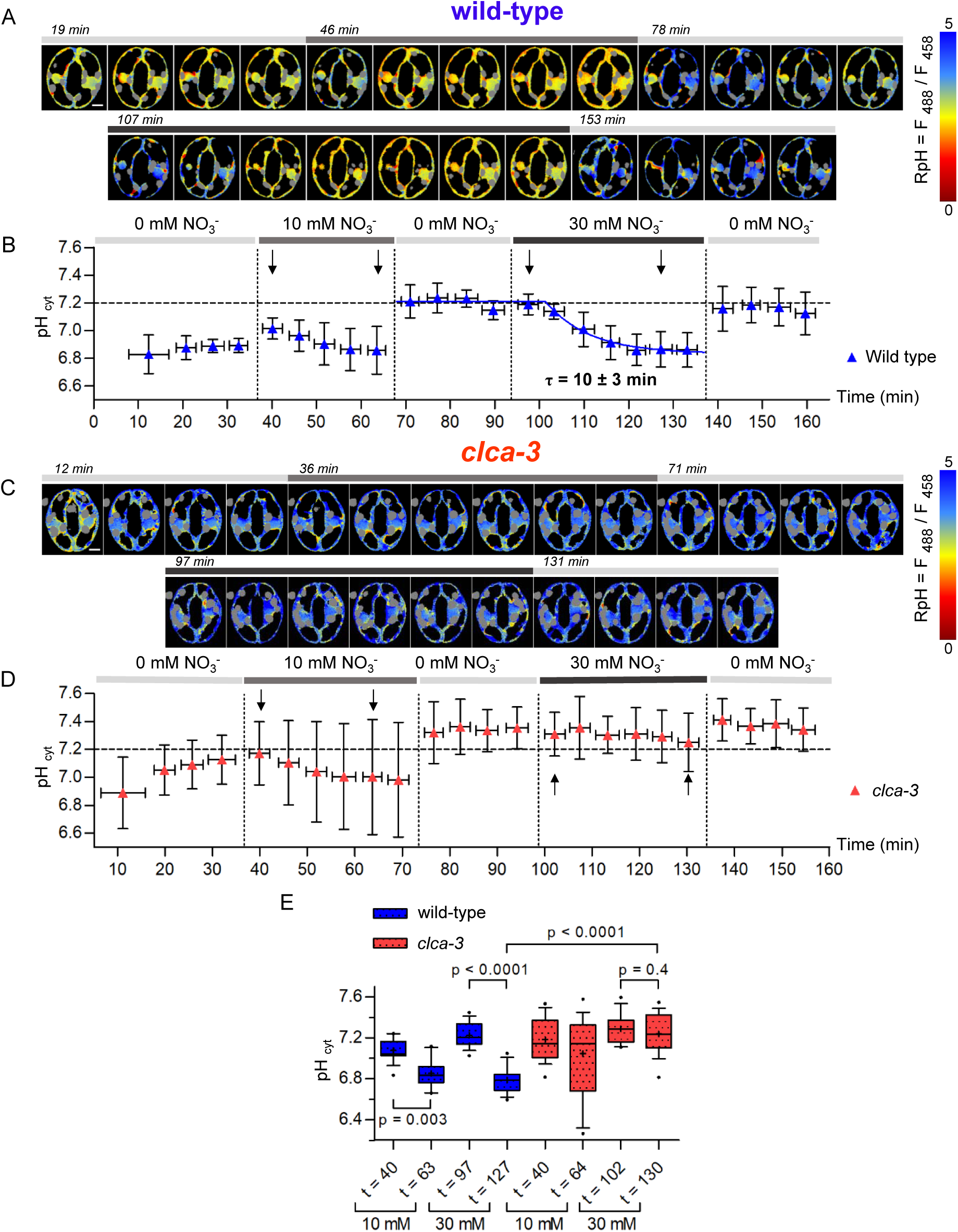
The vacuolar NO_3_^-^/H^+^ exchanger AtCLCa regulates pH_cyt_ in Arabidopsis stomata. Representative false colour ratio images of R_pH_ from wild type (A) and *clca-3* (C) stomata at different time points. Stomata were sequentially exposed to different extracellular conditions as indicated (top horizontal bar). Scale bar = 5 µm. Mean pH_cyt_ at each time point in (B) wild type (n=8) and (D) *clca-3* stomata (n=15). The pH_cyt_ depends on the applied extracellular concentration of KNO_3_ (top horizontal bar). Note the progressive decrease of pH_cyt_ after application of 30mM KNO_3_ in wild type (B) and the absence of pH_cyt_ changes in *clca-3* (D) stomata. The horizontal error bars show the time interval in which stomata were imaged. Vertical error bars represent standard deviation. Vertical dotted lines indicate changes of extracellular conditions. The horizontal dashed lines serves as reference for [anion]_cyt_ = 6 mM for comparison between Figure 2B, 4B and 4D. Grey areas in (A) and (B) show the position of the chloroplast and were excluded from the analysis. (E) box plots of the pH_cyt_ at different time points (black arrows in B and C). The data show that in wild-type stomata 30 minutes after exposure to 30 mM KNO_3_ the pH_cyt_ becomes significantly more acidic. Differently, in *clca-3* no significant acidification is observed. Blue boxes: wild type (n=17) ; red boxes: *clca-3* (n=15) stomata. Whiskers show the 10-90% percentile; the cross indicates the mean. The brackets indicate statistically significant differences.

Application of this 5-step protocol to wild type *pUBI10:ClopHensor* GCs demonstrated the dependency of [NO_3_^-^]_cyt_ on the applied extracellular KNO_3_ concentration, with [NO_3_^-^]_cyt_ being 1.64±0.32 and 4.74±1.52 mM in 10 and 30 mM KNO_3_, respectively (Fig. 3A, B,E). In presence of 10 mM KNO_3_ the [NO_3_^-^]_cyt_ leveled off in less than 4 minutes (Fig. 3B). Differently, in presence of 30 mM KNO_3_ in the extracellular medium the [NO_3_^-^]_cyt_ raised progressively with a time constant of τ=15±3 minutes (Fig. 3B). Interestingly, GCs maintained a [NO_3_^-^] gradient between the apoplast and the cytosol of about 6 fold when either 10 or 30 mM KNO_3_ was applied. In all cases, upon wash-out with NO_3_^-^-free medium, the [NO_3_^-^]_cyt_ dropped back to concentrations close to the limit of detection within 4 minutes. In *clca-3 pUBI10:ClopHensor* GCs the [NO_3_^-^]_cyt_ dynamics behaved similarly to wild type plants upon exposure to 10 mM KNO_3_ reaching 2.24±1.47 mM (Fig. 3C, D,E). Further, similarly to wild type, upon application of 30 mM KNO_3_ *clca-3 pUBI10:ClopHensor* GCs [NO_3_^-^]_cyt_ leveled-off at 6.34±2.91 mM (Fig. 3D). However, in contrast with wild type, [NO_3_^-^]_cyt_ raised faster, reaching a plateau in less than 3 minutes in *clca-3* compared to wild type *pUBI10:ClopHensor* GCs (Fig. 3D). These data are in agreement with the involvement of the AtCLCa exchanger in buffering cytosolic NO_3_^-^.

Furthermore, we found that the pH_cyt_ dynamics in wild type and *clca-3* were markedly different when extracellular KNO_3_ was applied (Fig. 4A-D). In wild type GCs, the pH_cyt_ stabilized at 6.89±0.05 at the beginning of the experiments. Then, exposure to 10 mM KNO_3_ induced an initial slight pH_cyt_ increase followed by a progressive and modest acidification of the cytosol. Washing-out with NO_3_^-^-free medium provoked a fast increase of the pH_cyt_ to 7.21±0.12. Then, upon perfusion with 30 mM KNO_3_, a progressive and marked acidification to 6.87±0.13 with a time constant of τ=10±3 minutes was observed. Finally, after washing-out in NO_3_^-^-free medium an alkalinisation to 7.16±0.16 was observed within 4 minutes. In *clca-3 pUBI10:ClopHensor* GCs, a modest pH_cyt_ acidification was observed upon exposure to 10 mM KNO_3_, as in wild type. However, such pH_cyt_ decrease was not statistically significant in *clca-3* plants (Fig. 4E). Remarkably, the perfusion of 30 mM KNO_3_, that induced a marked acidification in wild type GCs, did not induce any decrease of pH_cyt_ in *clca-3* GCs: pH_cyt_ remained stable at pH≈7.3 (Fig.4B, 4D). To exclude an effect of the KNO_3_ application sequence, we inverted step (2) and step (4) in the perfusion protocol and obtained the same results (Fig. S8). These findings clearly show that the presence of the 2NO_3_^-^/1H^+^ exchanger AtCLCa in the vacuolar membrane is associated with the pH_cyt_ modification detected in wild type GCs upon perfusion with 30 mM KNO_3_, suggesting a role of AtCLCa in the regulation of cytosolic pH.

A cross-reading of the above results strengthen the hypothesis of the role of AtCLCa activity in regulating cytosolic pH. In wild type plants, the dynamics of [NO_3_^-^]_cyt_ and pH_cyt_ were obviously related, as the recorded changes were in the same time scale, with correlated qualitative responses in the different experimental steps: when [NO_3_^-^]_cyt_ was stable, pH was stable; when [NO_3_^-^]_cyt_ was increasing, pH was decreasing (compare Fig. 3A, 3B with 4A, 4B). In contrast, in *clca-3 pUBI10:ClopHensor* GCs, [NO_3_^-^]_cyt_ changes were not mirrored by pH_cyt_ changes, showing that in *clca-3* the two processes were uncoupled (compare Fig. 3C, 3D with 4C, 4D).

## Discussion

The involvement of CLCs in severe genetic diseases in humans and their major physiological functions in plants has attracted considerable attention to these anion transport systems. Interestingly, the CLC family presents a dichotomy: the CLCs localized in the PM are anion channels, those localized in intracellular membranes are anion/H^+^ exchangers (1). A combination of electrophysiological, structural and biochemical data provided a detailed understanding of the mechanisms allowing the anion/H^+^ exchange or the anion channel behavior at a sub-molecular level in CLCs (1). However and despite intense research in the last decades, the cellular function of intracellular CLCs has remained elusive (1). So far, the role of intracellular CLCs was exclusively considered from the point of view of the organelle lumen, while the impact on the cytosolic compartment was overlooked. Nevertheless, when a CLC exchanger pumps anions into an organelle, it simultaneously releases a stoichiometric number of H^+^ in the cytosol. Therefore, intracellular CLCs have the capacity to influence cytosolic pH and regulate anionic homeostasis. To test this hypothesis *in vivo*, we used Arabidopsis GCs expressing the dual anion and pH biosensor ClopHensor to unravel the impact of a vacuolar CLC on the cytosol.

### ClopHensor is able to sense pH and Cl^-^ or NO_3_^-^ plant cells

ClopHensor is a genetically encoded biosensor originally developed in mammalian cells. Its photo-physical characteristics have been analyzed in depth (32, 36, 37). The advantageous properties of ClopHensor allow to measure, simultaneously and in the same cell, two important intracellular parameters, pH and the concentration of anions such as Cl^-^. Notably, changes in pH and [Cl^-^] can report the activity of different type of transmembrane transporters in the VM and PM of plant cells. Before using ClopHensor in plant cells we first checked its sensitivity towards other anions that, differently from animal cells, are present in the millimolar range in the cytosol (4) (Fig. 1). *In vitro* analysis demonstrated that ClopHensor is sensitive not only to Cl^-^ but also to NO_3_^-^, while it is insensitive to PO_4_^3-^, malate^2-^ and citrate^3-^ at the tested concentration (Fig 1). Furthermore, ClopHensor sensitivity to NO_3_^-^ is even higher than to Cl^-^. The analysis of the cytosolic [NO_3_^-^], [Cl^-^] and pH in living GCs demonstrated that ClopHensor is able to report dynamic changes of these parameters upon exposure to varying extracellular [NO_3_^-^] (Fig. 3-5). Interestingly, the cytosolic [NO_3_^-^] and [Cl^-^] we measured are in the same range as those previously reported in other cell types with selective microelectrodes (38–40). The agreement between our data and previous reports demonstrates the robustness of ClopHensor to measure [NO_3_^-^] and [Cl^-^] in Arabidopsis GCs. Concerning pH, ClopHensor displays a steep dynamic fitting cytosolic conditions (Fig. 1,2 and 4). The steepness of the pH sensitivity is particularly valuable to resolve subtle pH changes. Overall, our results demonstrate that ClopHensor can be used to measure [NO_3_^-^] and pH in GCs. Thus, ClopHensor is the first biosensor able to report [NO_3_^-^] in the cytosol; further it allows measuring pH in parallel. Given the link between anion and H^+^ transport in plant cells, this dual capacity is particularly relevant.

### A vacuolar CLC modifies cytosolic ion homeostasis

We conducted *in vivo* experiments with a conceptually simple design, yet never tested in plant cells. To reveal the impact of the activity of the vacuolar transporter AtCLCa, we challenged stomata with different extracellular media applied in a defined sequence (Fig. 2-4). Starting from an initial condition with no NO_3_^-^ or Cl^-^ in the extracellular medium and within the GCs, we applied different KNO_3_ and KCl based medium. In our conditions, cytosolic [Cl^-^] were below the sensitivity threshold of ClopHensor. We however obtained a first remarkable result: [NO_3_^-^]_cyt_ in GCs can undergo rapid variations (Fig. 2 and 3). To our knowledge, it is the first time that such modifications of [NO_3_^-^]_cyt_ are described. Former reports available from root epidermal cells or mesophyll protoplasts suggested that cytosolic NO_3_^-^ was, at least in the short term, stable (33, 40). These studies were using invasive approaches without challenging cells with different extracellular ion concentrations, possibly explaining why cytosolic NO_3_^-^ changes were not observed. Interestingly, our findings show that [NO_3_^-^]_cyt_ can change rapidly, within minutes (Fig. 2 and 3). This supports the hypothesis that cytosolic NO_3_^-^ variations may act as an intracellular signal. A role of cytosolic NO_3_^-^ to adjust cell responses to external nitrogen supply has been previously proposed (39, 41). A second remarkable observation we made is a progressive acidification of the cytosol in parallel with the cytosolic [NO_3_^-^] increase. Conversely, [NO_3_^-^]_cyt_ drop is paralleled by a rapid cytosolic pH increase (Fig 1). These findings clearly show a link between cytosolic NO_3_^-^ and pH changes, and suggests a common molecular mechanism underlying NO_3_^-^ and pH variations.

The detected changes in cytosolic pH and NO_3_^-^ integrate the transport reactions occurring at the PM and the VM as well as metabolic reactions and cytosolic buffer capacity. Our data suggest that the observed changes are due to H^+^ coupled transport reactions. In Arabidopsis cells, AtCLCa is the major H^+^ coupled NO_3_^-^ transporter in the vacuolar membrane (5, 29). Therefore, to test whether AtCLCa is responsible of the variations we detected in the cytosol, we conducted comparative experiments between GCs from wild type and from plants knocked-out for AtCLCa expressing ClopHensor (Fig. 3-4). We found that [NO_3_^-^]_cyt_ levels-off faster in *clca-3* GCs than in wild type when exposed to extracellular KNO_3_ (Fig 5). This proves that in *vivo* the vacuolar transporter AtCLCa buffers the cytosolic [NO_3_^-^], as expected from its function in accumulating NO_3_^-^ into the vacuole (5, 29). The most impressive consequence of knocking out AtCLCa was on the cytosolic pH (Fig. 4F). Indeed, in sharp contrast with wild type GCs, no pH acidification could be detected in *clca-3* GCs when cytosolic NO_3_^-^ increased. These unexpected finding reveals that AtCLCa solely accounts for the pH acidification we detected in wild type GCs. Of course, this does not mean that AtCLCa is the only H^+^ coupled NO_3_^-^ transport system operating in the PM and VM of GCs (Fig. 5). Rather, it indicates that upon increase in [NO_3_^-^]_cyt_ the transport activity of AtCLCa is high enough to overcome the pH buffering capacity of the cytosol. Therefore, the use of a biosensor like ClopHensor allowed us to detect *in vivo* the activity of an intracellular transporter, AtCLCa, and its impact on the intracellular ion homeostasis.

**Fig. 5.**
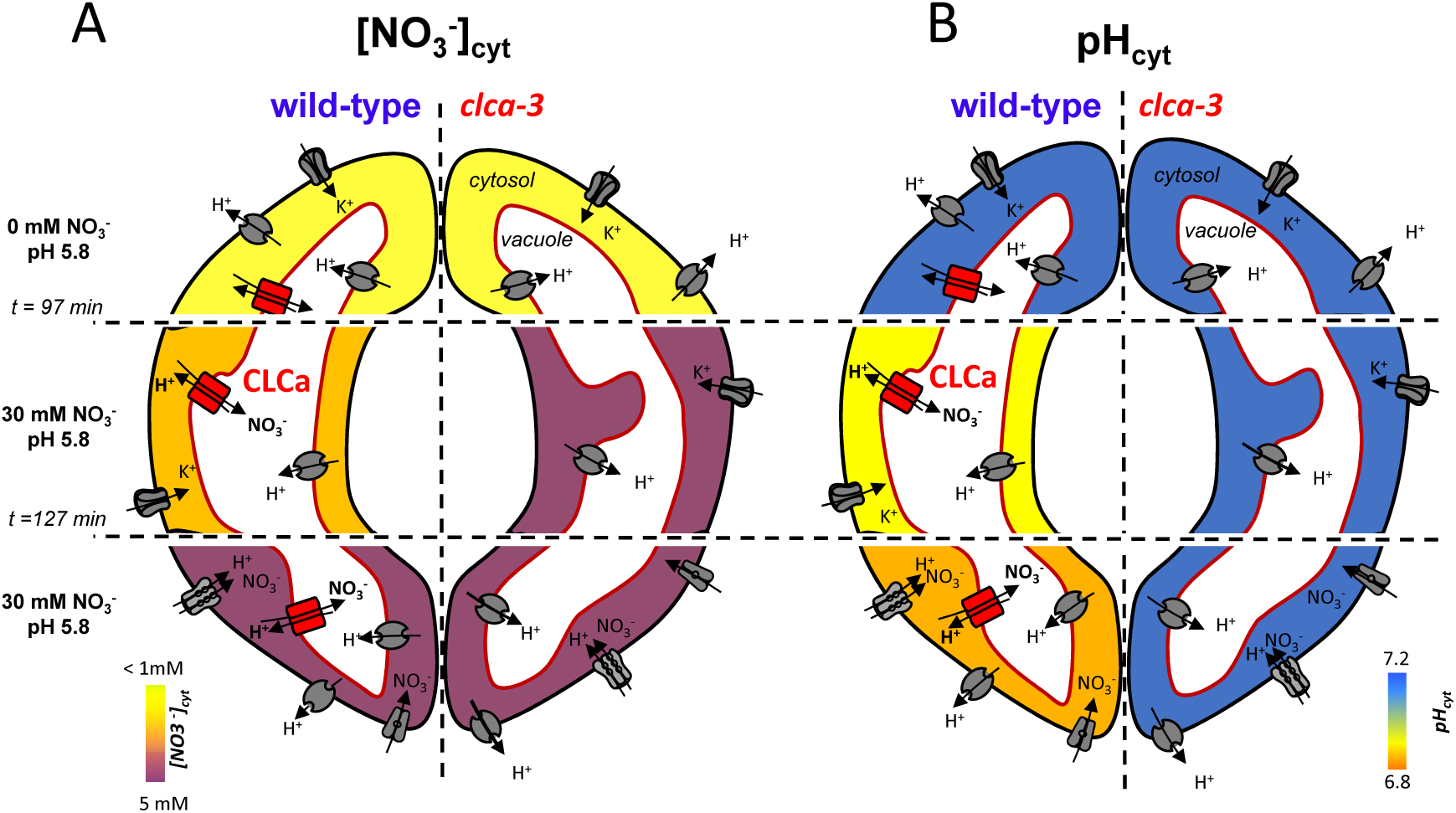
A vacuolar exchanger modifies cytosolic homeostasis in Arabidopsis stomata. (A) and (B), illustrations recapitulating the impact of the activity of AtCLCa on cytosolic NO_3_^-^ and pH homeostasis in guard cells (GCs). (A) In presence of 30 mM KNO_3_ in the extracellular media, NO_3_^-^ enters the cell via NO_3_^-^ transporters and channels residing in the plasma membrane. In wild type (*left GC)*, the vacuolar AtCLCa exchanger pumps NO_3_^-^ in the vacuole and consequently slows down the raise of [NO_3_^-^] in the cytosol. In the absence of AtCLCa (*right GC*), [NO_3_^-^] in the cytosol levels of in less than 4 minutes. (B) in presence of 30 mM KNO_3_ in the extracellular media the transport activity of AtCLCa releases H^+^ in the cytosol inducing an acidification in wild type GCs (*left GC*). In *clca-3*, in the absence of the exchanger activity, cytosolic acidification does not occur (*right GC*).

The finding that a vacuolar transporter influences cytosolic pH homeostasis is a major breakthrough in the understanding of the cellular functions of intracellular transporters. A potential role of H^+^ coupled transporters in the regulation of cytosolic pH was proposed in the ‘80s (42–44), but never demonstrated. Instead the role of intracellular ion transporters is nowadays commonly interpreted from “the point of view of the organelle”, focusing on how these transporters regulates ion homeostasis in the lumen of the organelles. Our data provide for the first time strong experimental evidence supporting the hypothesis that proton coupled intracellular transporters play an active role in the regulation of cytosolic pH. In the plant cell, the VM is commonly considered as a “second layer” with respect to the PM, which is postulated to rule over the intracellular conditions. Our findings clearly show that the effect of transporters in the VM is not “just to buffer the cytosol” to maintain cytosolic conditions at homeostatic values, but that they can actively modify the cytosolic conditions. AtCLCa is important in this process but it might not be the only one. It will be of interest to understand if and how other transporters like proton pumps or cation/H^+^ exchangers (e.g. NHX) as well as ion channels affect cytosolic homeostasis.

### Cytosolic pH control, a novel framework for CLC functions

The results presented here relate to a specialized plant cells type, the GCs. The effect of AtCLCa on cytosolic pH may account for the unexpected defect in stomata closure observed in *clca* knock-out plants, when its function in loading anions into the vacuole would rather lead to the prediction that it is solely involved in stomata opening (6). In this context, modification of cytosolic pH could be an important component of AtCLCa function, as cytosolic pH is an important parameter in cell signaling (45). Cytosolic pH modifications can modulate ion transport systems and enzymatic reactions to trigger stomata opening or closure. For example, the activity of vacuolar V-ATPase is modified by changes of the cytosolic pH conditions (46). Our finding also addresses the broader context of other eukaryotic CLC exchangers. Indeed, the function of intracellular CLCs has been interpreted assuming that their only role was to regulate the lysosomal, endosomal or vacuolar lumen conditions (1). However, in the case of the lysosomal CLC-7 and endosomal CLC-5 their cellular function remains unclear. CLC-7 was proposed to acidify the lysosomal lumen, but only modest and controversial effects were detected (13, 15). In the case of CLC-5 endosomes from knock-out mice present impaired luminal acidification (31). Nonetheless, the connection between endosomal acidification and the severe defects caused by CLC-5 mutations in Dent’s disease is still unclear (1). Indeed, renal failure associated with some mutations in CLC-5 present impaired endocytosis in tubular cells which is independent of endosomal acidification (31). Intriguingly, cytosolic pH is known to affect endocytosis (47, 48). The findings we report here suggest that in eukaryotic cells intracellular CLCs are part of the cytosolic pH balance machinery. This opens a novel perspective on the function of these exchangers in eukaryotic cells and may provide a novel framework to understand the pathophysiological disorders caused by mutations in human CLC genes.

## Methods

### Protein expression and purification for *in situ* calibration of ClopHensor

Recombinant ClopHensor was expressed with a strep-tag in *Escherichia coli* (ΔH5α strain) with *pIBA2:ClopHensor* vector. Expression was induced with tetracycline at 25°C for 13 hours. Bacteria were lysed by sonication with extraction buffer (100mM (NH_4_)_2_SO_4_; 50mM Tris-H_2_SO_4_ pH 7.5; 5mM β-mercapto-ethanol; 1mM PMSF and a protease inhibitor (cOmplete™), EDTA-free Protease Inhibitor Cocktail, Sigma Aldrich). ClopHensor was bound to Strep-Tactin Sepharose 50% suspension beads (IBA lifescience). Beads were equilibrated (100mM Tris-H_2_SO_4_ (pH 8), 50mM (NH_4_)_2_SO_4_, 1mM EDTA) at 4°C. Protein extracts were added to equilibrated beads overnight. For confocal imaging, the beads were washed 5 times in control buffer (Supplementary Information) at room temperature. NaCl or NaNO_3_ were added to the control buffer to 0; 1; 10; 30; 100 and 300mM.

### Cloning and molecular biology

ClopHensor was cloned in the pENTR2B (ThermoFischer) vector with KpnI and XhoI restriction sites (primers in Table S2). It was then sub-cloned in the homemade plant expression vector pMubi32. pMUBI32 is the backbone of pMDC32 (49) where the 35S promoter was substituted by the ubiquitin 10 promoter. *pUBI10:ClopHensor* was introduced in *Agrobacterium tumefaciens* (GV3101 strain) for stable Arabidopsis transformation.

### Plant material

To generate *Arabidopsis* lines stably expressing ClopHensor, Arabidopsis (Col-0 ecotype) were transformed by the floral dip method (50) with *Agrobacterium tumefaciens* strains carrying the *pMUBI32:ClopHensor* plasmid. Transgenic Arabidopsis expressing *pUBI10:ClopHensor* were selected on agar plates containing ½ MS medium containing 15µg/l of hygromycin B. Plants from homozygous T_3_ and T_4_ generations were used for experiments. To obtain *clca-3 pUBI10:ClopHensor* plants, we crossed wild type *pUBI10:ClopHensor* plants with *clca-3* knock out (Gabi Kat line GK-624E03-022319). Homozygosity of the mutants was tested by PCR (primers in Table S2). Plants used for imaging were grown *in vitro* (Supplementary Information) in culture chambers (Sanyo MLR350) at 21°C with a 16h photoperiod (60 µmol photon.m^-2^.s^-1^) for 14 days.

### Confocal Laser Scanning Microscopy

12 bits images were acquired with a Leica SP8 upright confocal laser scanning microscope (CLSM), equipped with a 63x Plan Apochromat oil-immersion objective (NA= 1.4). Sequential excitation mode (561, 488 and 458 nm) was used to collect images on GaAsP Hybrid photon detectors using the photon counting mode (Hamamatsu). ClopHensor Emission was detected at 500-550nm upon excitation at 458nm and 488nm (Argon laser), and at 600-625nm upon excitation at 561nm (DPSS laser). Chloroplasts were excited at 488 nm and collected at 600-625nm. Transmitted light images were obtained with a PMT detector. The whole system was driven by LAS X software (Leica). Images were analyzed by ImageJ with homemade macro as described in Fig. S5.

### Image analysis

Detailed description of the image analysis procedure is described in Fig. S5. Briefly, on each fluorescence image the background was subtracted. To compensate laser intensity fluctuations and cell to cell variability, fluorescence images were corrected with mean intensity of the corresponding transmission image. The region of interest, corresponding to the cytosolic compartment and nucleoplasm were selected applying a manual threshold (> 100 photons/pixel) to the at 458 nm excitation image. Chloroplasts were selected by automatic threshold (*Otsu Dark* method) and subtracted for the analysis. The ratio images for RpH (F_488_/F_458_) and R_Anion_ (F_458_/F_561_) were obtained by dividing the fluorescence images. The mean <RpH> or <R_Anion_> were extracted from the ratio images. The pH and anion concentration values were calculated from the <RpH> or <R_Anion_> as in (32) (Supplementary Information)

### Sample preparation and perfusion

For stomata measurements, leaves of 14 days olds plantlets grown *in vitro* were glued (Adapt™ Medical Adhesive) on a microscope coverslip, and peeled to isolate the abaxial epidermis (20). The coverslip was mounted on a home-made perfusion chamber with a volume of 100-200 µl of NO_3_ free-medium (Supplementary Information). For *in vivo* pH titration stomata were perfused with NH_4_-Acetate buffers at the indicated pHs (Supplementary Information). Every KNO_3_ or KCl buffer was prepared from 0 NO_3_ buffer and supplemented with corresponding quantity of KNO_3_ or KCl. The pH was adjusted to 5.8 with KOH. In all media the osmolarity was set at 100 mosmol with sorbitol.

### Statistical analyses

Statistical analysis was performed with Prism software. Data were analyzed with two-tailed non parametric Mann Whitney test and the P value is presented for each pair tested.

## Acknowledgments

This work was supported by LabEx Saclay Plant Sciences-SPS (ANR-10-LABX-0040-SPS). This work has benefited from the facilities and the expertise of Imagerie-Gif microscopy platform, which is supported by France-BioImaging (ANR-INBS-04, ‘Investments for the future’), and by ‘Saclay Plant Science’ (ANR-11 IDEX-0003-02). We would like to thanks D. Arosio (CNR, Trento, Italy) for providing plasmid with ClopHensor and for discussion, R. Le Bars (Imagerie-Gif) for help, M. Bianchi (I2BC, Gif sur Yvette) and S. Filleur (I2BC, Gif sur Yvette) for discussion.

## Authors Contribution

A.D.A, S.T. conceived and designed research. E.D., L.B, A.D.A performed experiment and analysed data. L. B and A.D.A. developed macro for the analysis. A.D.A, S.T, B.S.J, wrote the manuscript. E.D. and L. B. prepared Figures.

