## Supplementray Information for "Dynamic measurement of cytosolic pH and [NO_3_^-^] uncovers the role of the vacuolar transporter AtCLCa in the control of cytosolic pH"

This pdf contains :

Supplementary text

Figure S1 to S7

Tables S1 and S2

### Supplementary text:

#### *In vitro* calibration, data analysis

*In vitro* pH titration data (Fig 1E) were fitted with equation (S1) like in Arosio et al. 2010 (1).

$$R_{pH} = \frac{\mathbf{R_B} + \mathbf{R_A}\, 10^{(pKa - pH)}}{1 + 10^{(pKa - pH)}} \qquad \text{(S1)}$$

$[NO_3^-]$ or $[Cl^-]$ titration of $R_{Anion}^{norm}$ data were fitted according to equation (S2), where C is the maximum quenching of ClopHensor induced by the anion.

$$R_{Anion} = 1 - \frac{C}{1 + 10^{(pKa - pH)}} \qquad \text{(S2)}$$

#### Calculation of $[NO_3^-]$, $[Cl^-]$ and pH in stomata

Calculation of cytosolic pH and anion was performed as previously described by Arosio et al. (2010)(1); Briefly, in each stomata $pH_{cyt}$ was calculated using equation (3). We used pKa value of 6.9.

$$pH = \mathbf{pKa} + \log\left(\frac{R_{pH} - \mathbf{R_A}}{\mathbf{R_B} - R_{pH}}\right) \qquad \text{(S3)}$$

The $R_A$ and $R_B$ values used were 0.5 and 4.9, respectively. For each stomata, the RpH was calculated as the mean over the ratio all pixels of the ratio between the fluorescence detected at 488 and 458 nm excitation wavelengths.

Once the cytosolic pH calculated, we derived the corresponding dissociation ($Kd_{Anion}(pH)$) constant of ClopHensor at each pH with equation (S4), where $Kd_{anion}$ is the limit dissociation constant of $NO_3^-$ or $Cl^-$ at acidic pH when ClopHensor is in the protonated form (1).

$$Kd_{Anion}(pH) = \mathbf{Kd_{Anion}}\left(\frac{1 + 10^{(\mathbf{pKa} - pH)}}{10^{(\mathbf{pKa} - pH)}}\right) \qquad \text{(S4)}$$

To calculate the $[NO_3^-]$ or $[Cl^-]$ in the cytosol of each stomata we used equation (S5). Here, $R_{Anion}$ is the mean over all pixels of the ratio between the fluorescence detected after excitation at 458 nm and the fluorescence detected after excitation at 561 nm. The $R^0_{Anion}$ is the value of $R_{Anion}$ in the absence of $NO_3^-$ or $Cl^-$, calculated the $R^0_{Anion}$ for each stomata in $NO_3$ free buffer at the beginning of each experiment.

$$[Anion] = Kd_{Anion}(pH)\left(\frac{R^0_{Anion} - R_{Anion}}{R_{Anion}}\right) \quad (S5)$$

In all cases, the fluorescence detected at excitation wavelengths was normalised with the mean intensity of the corresponding transmission image (Fig S6C).

In figures 3 and 4 to estimate the time constant of the cytosolic $[NO_3^-]$ increase data were fitted with an exponential function preceded by a plateau (equation S6). In equation (6), $<[NO_3^-]_0>$ is the mean value of $[NO_3^-]$ in 0 $NO_3$ media before the application of 30 mM $KNO_{3,extracell}$, $\tau$ is the time constant of the raise (min) and t is the time in minutes, $t_0$ is the time when the extracellular $KNO_3$ was applied. The plateau is the maximum $[NO_3^-]_{cyt}$ reached in presence of 30 mM $KNO_{3,extracell}$.

$$[NO_3^-] = <[NO_3^-]_0> + (plateau - <[NO_3^-]_0>)\,(1 - exp^{(-(t - t_0)/\tau)}) \quad (S6)$$

The pH values were fitted with a decay function preceded by a plateau (equation 7) to estimate the time constant of the decrease ($\tau$):

$$pH = plateau + (<pH_0> - plateau)\,exp^{(-(t - t_0)/\tau)} \quad (S7)$$

$<pH_0>$ is the mean pH value measure in $NO_3$ free conditions before application of 30 mM $KNO_{3,extracell}$. The plateau is the minimum $pH_{cyt}$ reached in presence 30 mM $KNO_{3,extracell}$.

**Colorimetric dosage of $NO_3^-$and $Cl^-$ contents in plants**

Plants were grown *in vitro* on agar plates in a $NO_3^-$ free medium were collected at different days after germination (DAG Supplementary Table 1). For each batch of plants 20 to 60 seedlings were weighted and frozen in liquid nitrogen. Subsequently plants were grinded with steel beads by a mixer mill (Retsch MM400) for 1min at 24 Hz. The obtained powder was dissolved in 500µl of $ddH_2O$. After 3 successive cycles of freezing-defrosting, samples were centrifuged at 13 000rpm for 5min to eliminate cell debris and collect the supernatant. $NO_3^-$ content was determined as described in Wege et al. 2014 (2) by colorimetric dosage using a reactive solution (500mM HCl; 16mM $VCl_3$; 0.2mM N-1-naphtylethylenediamine; 6mM sulfamilamide (3). 100µl of the reactive solution was added to 100µl of the sample. The absorbance at 540nm was measured after 1 hour of incubation at room temperature with a spectrophotometer. $Cl^-$ content was determined by colorimetric assay using the Ferricyanide method (4).

**DNA extraction and PCR blot**

DNA was extracted from a $4mm^2$ of leaf tissue from 3 weeks after germination. Samples were frozen in liquid nitrogen, then grinded with steel beads by a mixer mill (Retsch MM400) for 1min at 24Hz. Samples were dissolved in an extraction solution (200mM Tris-HCl (pH 7.5); 250mM NaCl; 250mM EDTA and 0.5% SDS). After centrifugation at 13 700rpm for 10min, addition of isopropanol to the supernatant enabled DNA to precipitate. DNA was collected by centrifugation at 13 700rpm for 10min and washed with ethanol (70%). Once the ethanol completely evaporated, DNA was solubilized in $ddH_2O$. PCRs was performed with primers in Table S2.

**Electrophysiological recordings**

Patch clamp recordings were performed in whole vacuole obtained from on 4-5 weeks old plant leaves as described in (5). Currents were evoked in response to 5s pulses from -77 to +43mV in +20mV increments followed by 3 seconds of tail pulse at -67mV and with a holding potential at -17mV (liquid junction potential corrected off line). Bis-Tris-Propane (BTP) was used as an impermeable cation to minimize cationic currents across the tonoplast. The bath solution (cytosol) contained 7.5mM BTP; 15mM HCl; 0.1mM $CaCl_2$; 2mM $MgCl_2$; 15mM MES at pH 7 and an osmolarity of 640mOsm. The pipette solution (vacuole) contained 100mM BTP; 200mM $HNO_3$; 1mM $CaCl_2$; 5mM $MgCl_2$; 5mM MES at pH 5.5 and an osmolarity of 590mOsm. The osmolarity of solutions was adjusted by adding sorbitol. Recordings were performed using an EPC10 amplifier (HEKA ElectroniK) controlled with Patchmaster software (HEKA ElectroniK). Data analysis was done with Fitmaster (HEKA ElectroniK).

**Buffers:**

*Sepharose Beads control buffer* 60mM $(NH_4)_2SO_4$; 10mM MES (pH 5.5; 6) or BTP (pH 7; 7.5; 8; 9) or CAPS (pH 10), pH was adjusted with $H_2SO_4$ or NaOH.

*$NH_4$-acetate buffer:* 50mM $NH_4$-acetate; 0.1mM MgSO4; 0.1mM CaOH and 10mM citric acid (pH 5) or MES (pH 5.5; 6.5) or BTP (pH 7; 7.5; 8.5; 9), pH was adjusted with $H_2SO_4$ or NaOH.

*$NO_3$ free culture medium:* 1mM $CaSO_4$; 1mM $KH_2PO_4$; 500µM $MgSO_4$; 50µM NaFeEDTA; 50µM $H_3BO_3$; 12µM $MnCl_2$; 1µM $ZnSO_4$; 1µM $CuSO_4$; 30nM $(NH_4)_6Mo_7O_{24}$; 10g/l saccharose; 0.5g/l MES; 2mM Glutamine; 0.5mM $K_2SO_4$; 0.8% of phytagel. The pH was adjusted to 5.8 with KOH.

*$NO_3$ free medium:* 1mM $CaSO_4$; 1mM $KH_2PO_4$; 500µM $MgSO_4$; 50µM NaFeEDTA; 50µM $H_3BO_3$; 12µM $MnCl_2$; 1µM $ZnSO_4$; 1µM $CuSO_4$; 30nM $(NH_4)_6Mo_7O_{24}$; 10g/l saccharose; 0.5g/l MES; 2mM Glutamine; 0.5mM $K_2SO_4$. The pH was adjusted to 5.8 with KOH.

### Supplementary Figures:

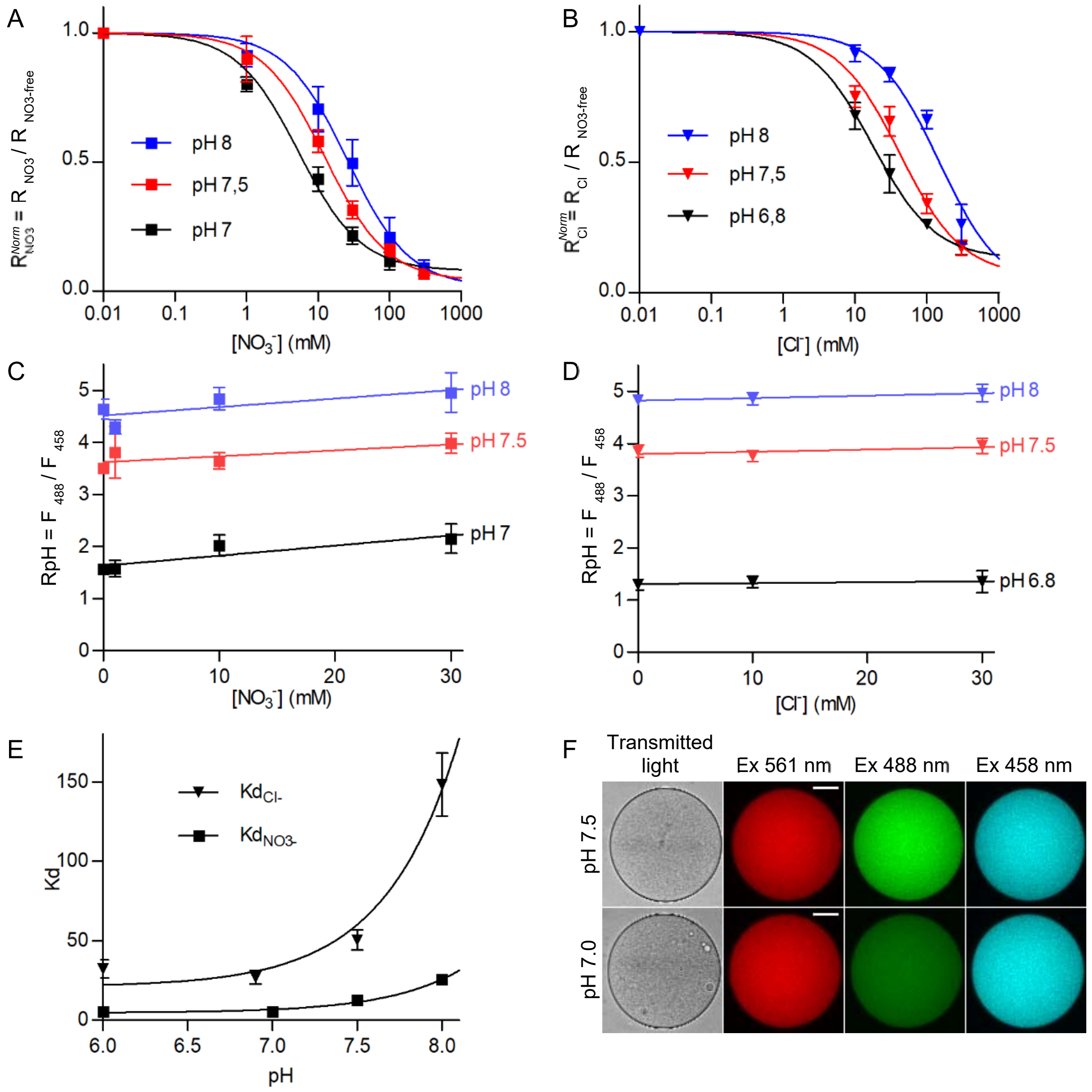


**Fig. S1. *In vitro* analysis of ClopHensor sensitivity towards $NO_3^-$, $Cl^-$ and pH.** (A , B) ClopHensor dose-response analysis for $NO_3^-$ (A) and $Cl^-$ (B). Different concentrations (0; 1; 10; 30; 100; 300 mM; n≥15 beads for each condition) of $NaNO_3$ (A) and NaCl (B) were tested at different pH, 8; 7.5; 7 (A); 6.8 (B). Data were normalized in respect of control condition value and fitted with equation S2. (C , D) Analysis of the influence of $NO_3^-$ and $Cl^-$ concentration on RpH measurements. Plot of RpH at pH 8 (blue), 7.5 (red) and 7; 6.8 (black) in presence of $NaNO_3$ (C) or NaCl (D) (0; 1; 10; 30 mM; n=15 beads for each condition). Data were fitted with a linear equation. (C) pH dependency of ClopHensor's affinity towards nitrate or chloride. $K_d$ measurements with $NaNO_3$ (squares) and NaCl (triangles) at different pH 6; 7; 7.5 and 8. Data were fitted with equation S2. (F) Confocal images of sepharose beads binding purified ClopHensor at pH 7.5 (top) and pH 7.0 (bottom) in control conditions. Excited at 561, 488 and 458 nm wavelengths. Scale bar = 30 µm. Error bars are the standard deviations.

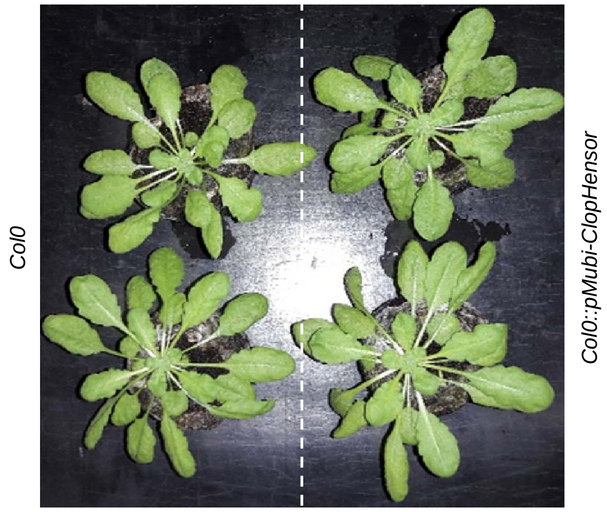


**Fig. S2. The expression of ClopHensor does not affect Arabidopsis development.** Arabidopsis plants from *Col-0* plants (*right*) and Col-0 plants expressing *pUBI10:ClopHensor* (left) grown in culture chambers with the same conditions with a 16 hours photoperiod.

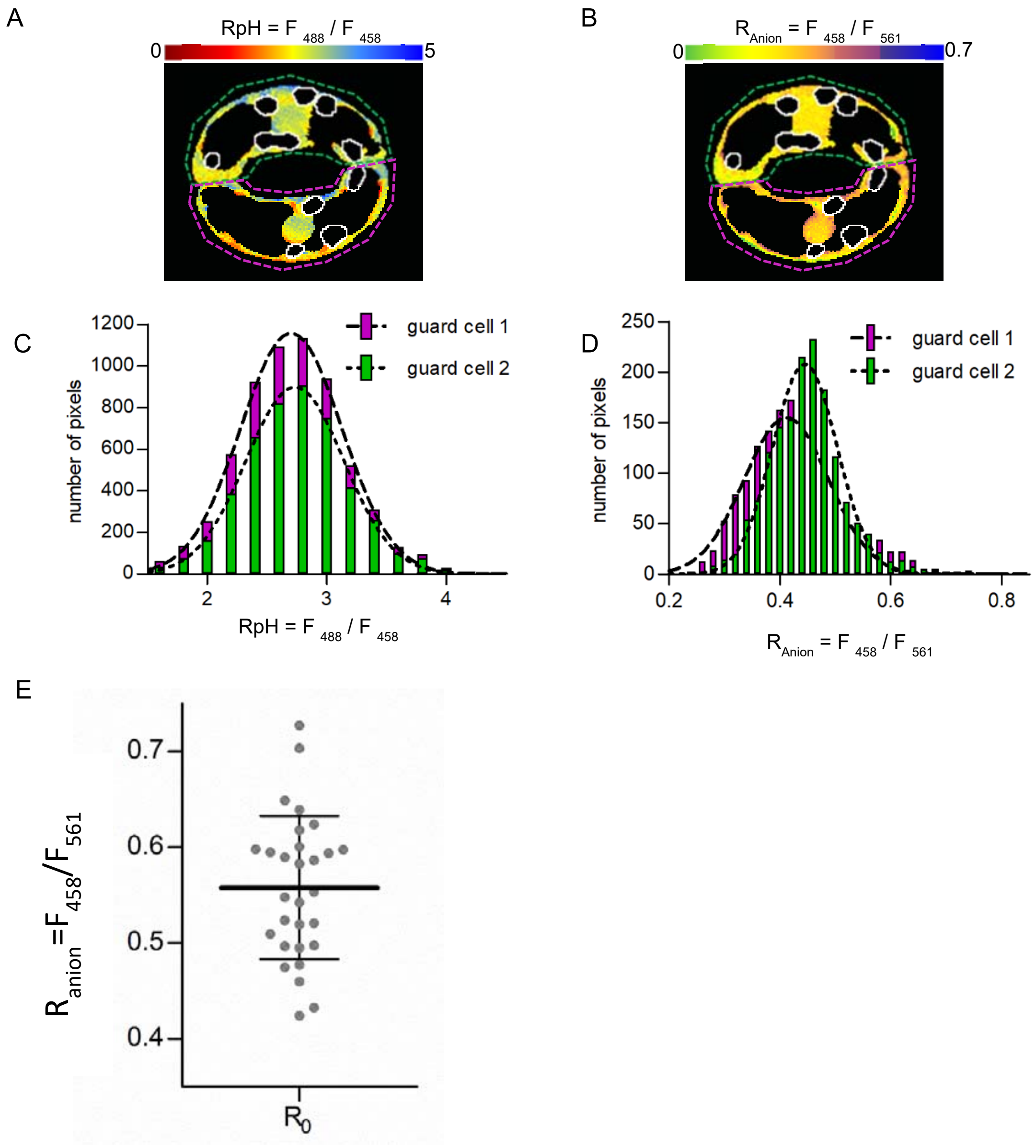


**Fig. S3. The two GCs of a stomata display overlapping RpH and $R_{Anion}$ distributions.** (A, B) False colour ratiometric images of a stomata from wild type plant expressing ClopHensor illustrating the RpH (A) and $R_{Anion}$ (B) in the cytosolic compartments. White contours illustrate the localization of the chloroplasts that are subtracted for the analysis. (C, D) distribution histograms of RpH (C) and $R_{Anion}$ (D) values from each pixels from images in A and B. The two GCs are here analysed individually. The data show that the RpH and $R_{anion}$ of the two individual GCs overlap. (E) $R_{Anion}$ calculated in stomata obtained from plants grown in $NO_3^-$free medium 14 DAG and exposed to extracellular $NO_3^-$free medium during imaging. Each point represents the mean $R_{Anion}$ of a stomata (n=29). Bars represent mean and standard deviation.

**wild-type**

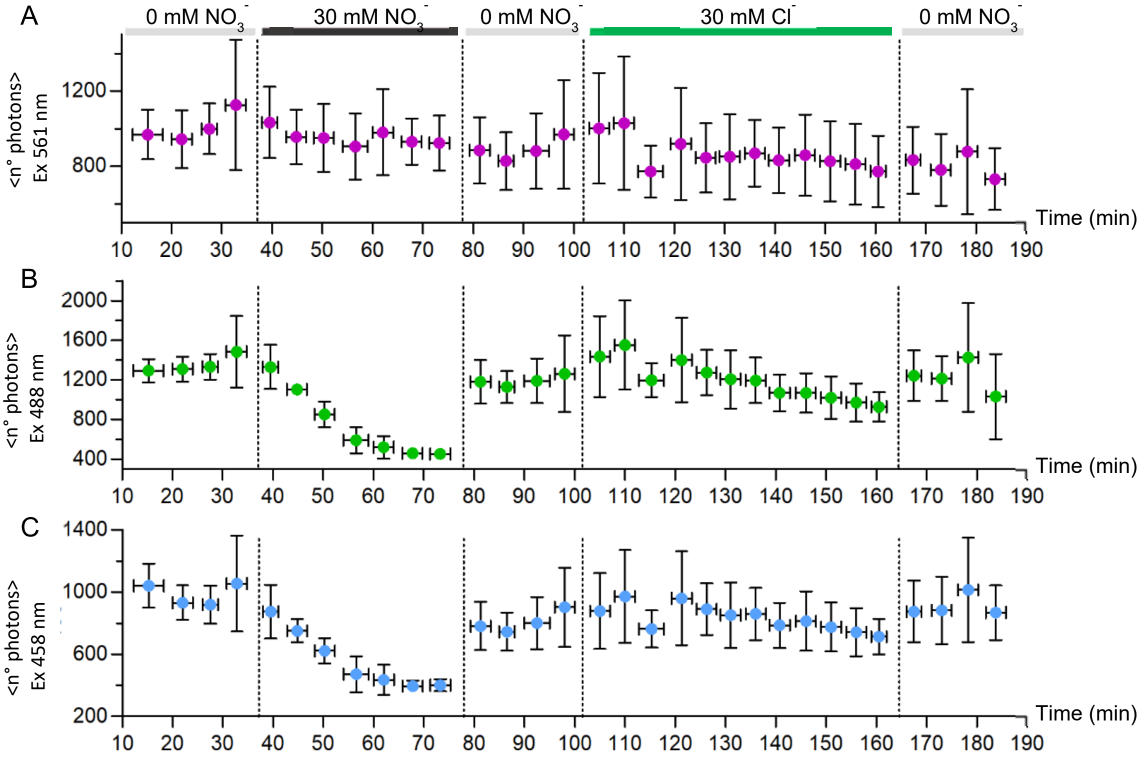


**Fig. S4. Mean number of photons detected during an extracellular buffer exchange in wild type *pUBI10:ClopHensor* stomata.** Mean number of photons emitted after excitation of ClopHensor at 561 (A), 488 (B) and 458 nm (C) measured during the experiment presented in Fig.2. A, mean number of photons detected after excitation at 561 nm of the DsRed moiety of ClopHensor. The data show that the number of photon detected is stable during the experiment, indicating that the amount of the biosensor is constant over the experiment. B and C, mean number of photons detected after excitation at 488 nm (B) and 458 nm (C) shows that upon the addition of 30 mM $KNO_3$ the $E^2GFP$ moiety of ClopHensor is reversibly quenched. The mean number of photon detected at the beginning and at the end of the experiment are not significantly different indicating that under the used conditions the $E^2GFP$ moiety of ClopHensor is not subject to photo bleaching.

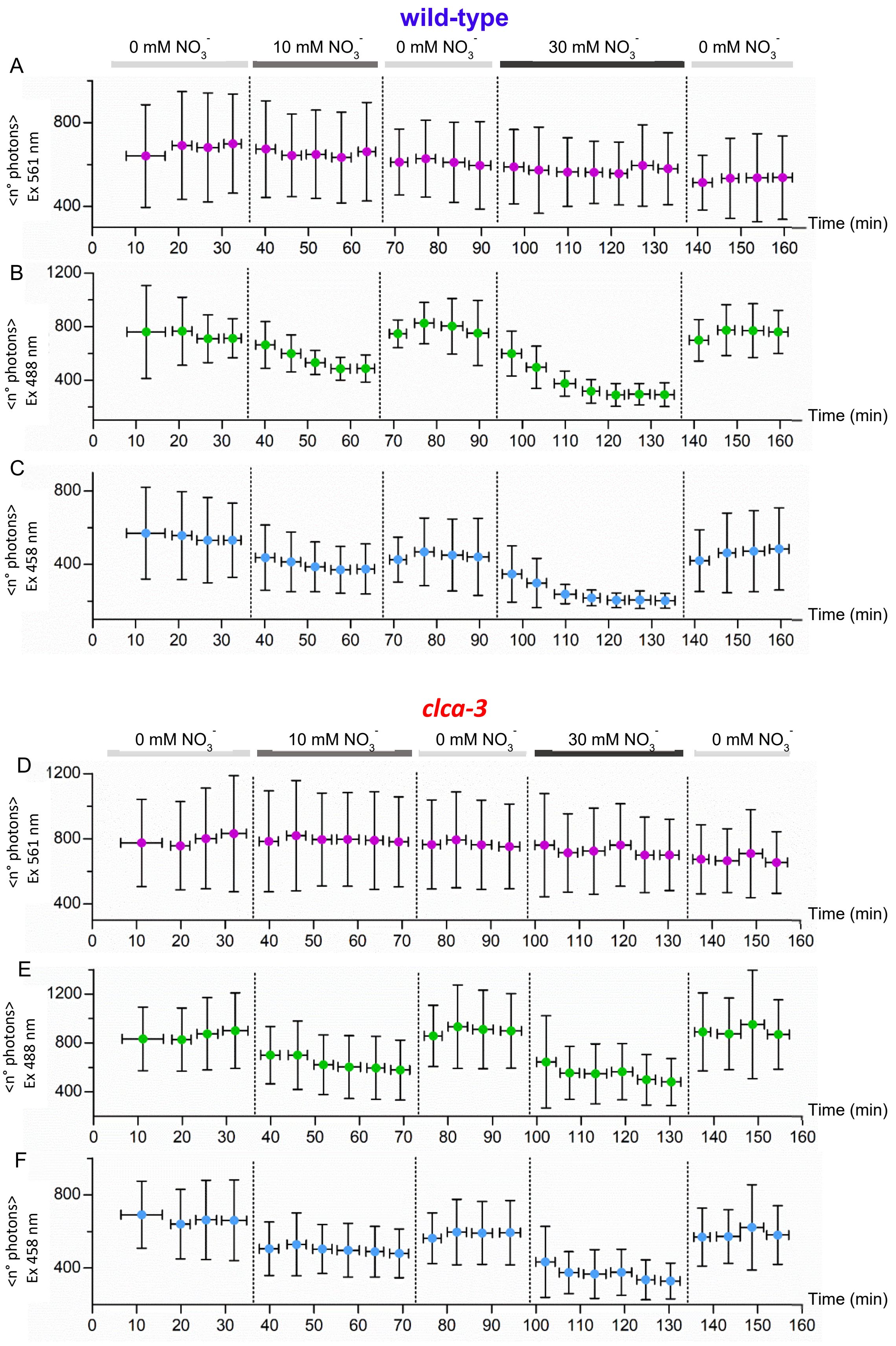


**Fig. S5. ClopHensor is not subject to photo bleaching in both wild type *pUBI10:ClopHensor* and *clca-3 pUBI10:ClopHensor* GCs.** Analysis of the mean number of photons detected over time in wild type *pUBI10:ClopHensor* stomata after excitation of ClopHensor at 561 (A), 488 (B) and 458 nm (C) wavelengths. The data correspond to Fig 3 and 4 (n=8). Analysis of the mean number of photons detected over time in in *clca-3 pUBI10:ClopHensor* stomata after excitation at 561 (D), 488 (E) and 458 nm (F) wavelengths corresponding to Fig 3 and 4 (n=15). The mean number of photons detected at the beginning and at the end of the experiment in both genotypes are stable indicating that under the used conditions ClopHensor is not affected by photo bleaching.

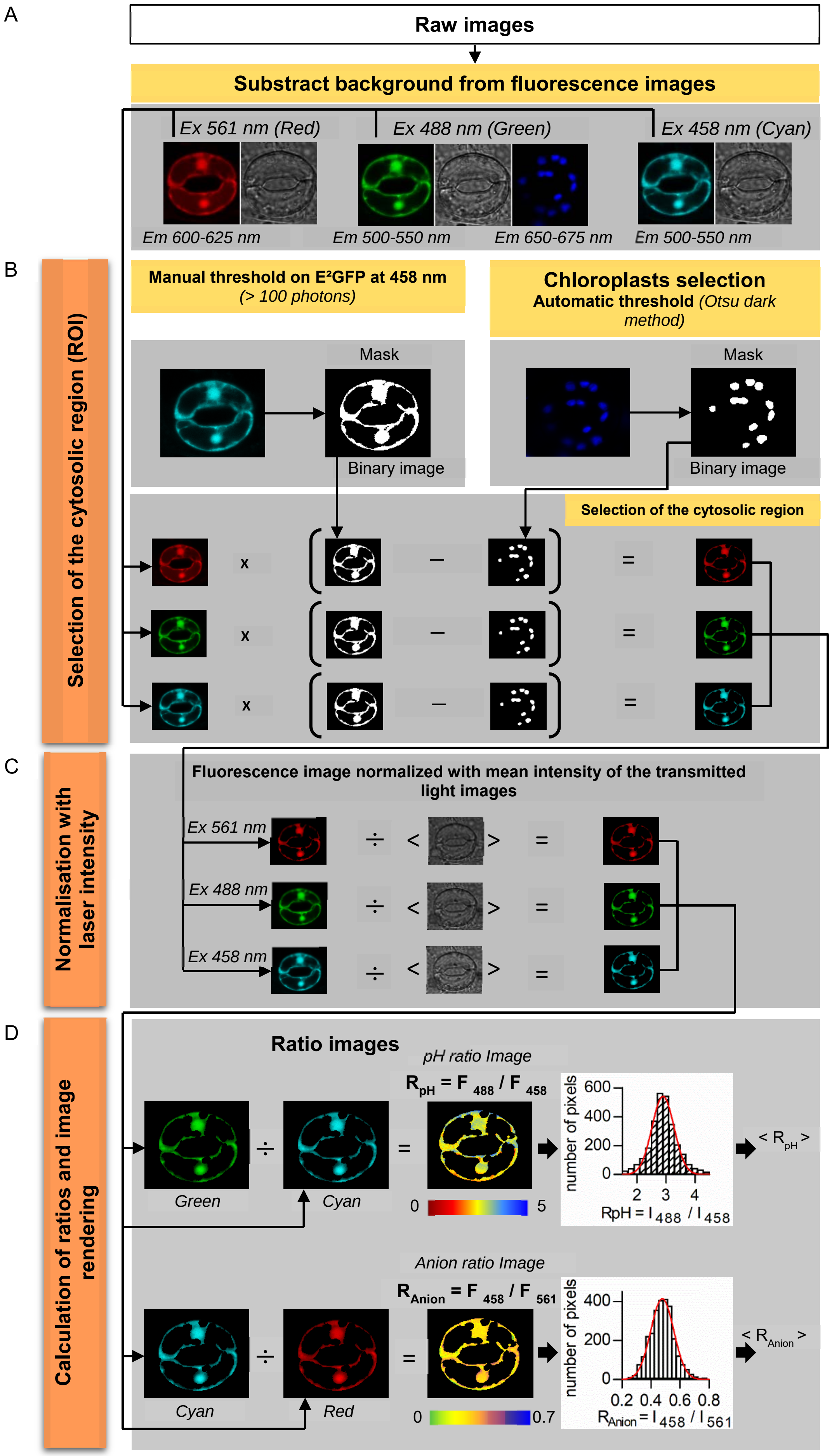
A
Raw images
Substract background from fluorescence images
Ex 561 nm (Red)
Ex 488 nm (Green)
Ex 458 nm (Cyan)
Em 600-625 nm
Em 500-550 nm
Em 650-675 nm
Em 500-550 nm
B
Selection of the cytosolic region (ROI)
Manual threshold on E²GFP at 458 nm
(> 100 photons)
Chloroplasts selection
Automatic threshold (Otsu dark method)
Mask
Binary image
Mask
Binary image
Selection of the cytosolic region
C
Normalisation with laser intensity
Fluorescence image normalized with mean intensity of the transmitted light images
Ex 561 nm
Ex 488 nm
Ex 458 nm
D
Calculation of ratios and image rendering
Ratio images
pH ratio Image
R pH = F 488 / F 458
Green
Cyan
0
5
number of pixels
600
400
200
0
2 3 4
RpH = I 488 / I 458
< R pH >
Anion ratio Image
R Anion = F 458 / F 561
Cyan
Red
0
0.7
number of pixels
400
200
0
0.2 0.4 0.6 0.8
R Anion = I 458 / I 561
< R Anion >

**Fig. S6. Image analysis workflow.**

Panels illustrate the image analysis procedure applied to the confocal images of stomata. The whole work flow was organized in an *in house* Image J macro.

**(A),** confocal images obtained exciting at the 3 wavelengths ClopHensor, 561 (Em 600-625), 488 (Em 500-550; 650-675 nm for chlorophyll signal) and 458 nm (Em 500-550 nm). The background noise was estimated averaging an area without cell in each fluorescence image and was then subtracted.

**(B),** selection of the GCs cytosolic region (ROI). A cytosolic mask was generated using a manual threshold based on the number of detected photons (>100) to limit the effects of shot noise (6). The threshold was performed on the cyan image (Ex 458 nm). A second mask to select the chloroplasts was obtained with an automatic threshold (Otsu dark method). These masks were then applied to each fluorescent image to isolate the signal of the cytosolic and nucleoplasmic region, thus removing the chloroplasts from the analysis.

**(C),** once the cytosol was selected each fluorescent image was normalized to compensate for laser intensity fluctuations. Every image from panel B was divided by the mean intensity that was simultaneously measured on the corresponding transmitted light image. The mean intensity of the transmitted light image is a reliable measure of the intensity of laser used to excite ClopHensor.

**(D)**, calculations to obtain ratio images and the mean RpH and $R_{Anion}$. The different fluorescence images were divided to obtain a ratio image of RpH (F488/F458) and $R_{Anion}$ (F458/F561). The average ratio was directly computed from these images. The distribution RpH and $R_{Anion}$ in the ratio images follows a Gaussian distribution.

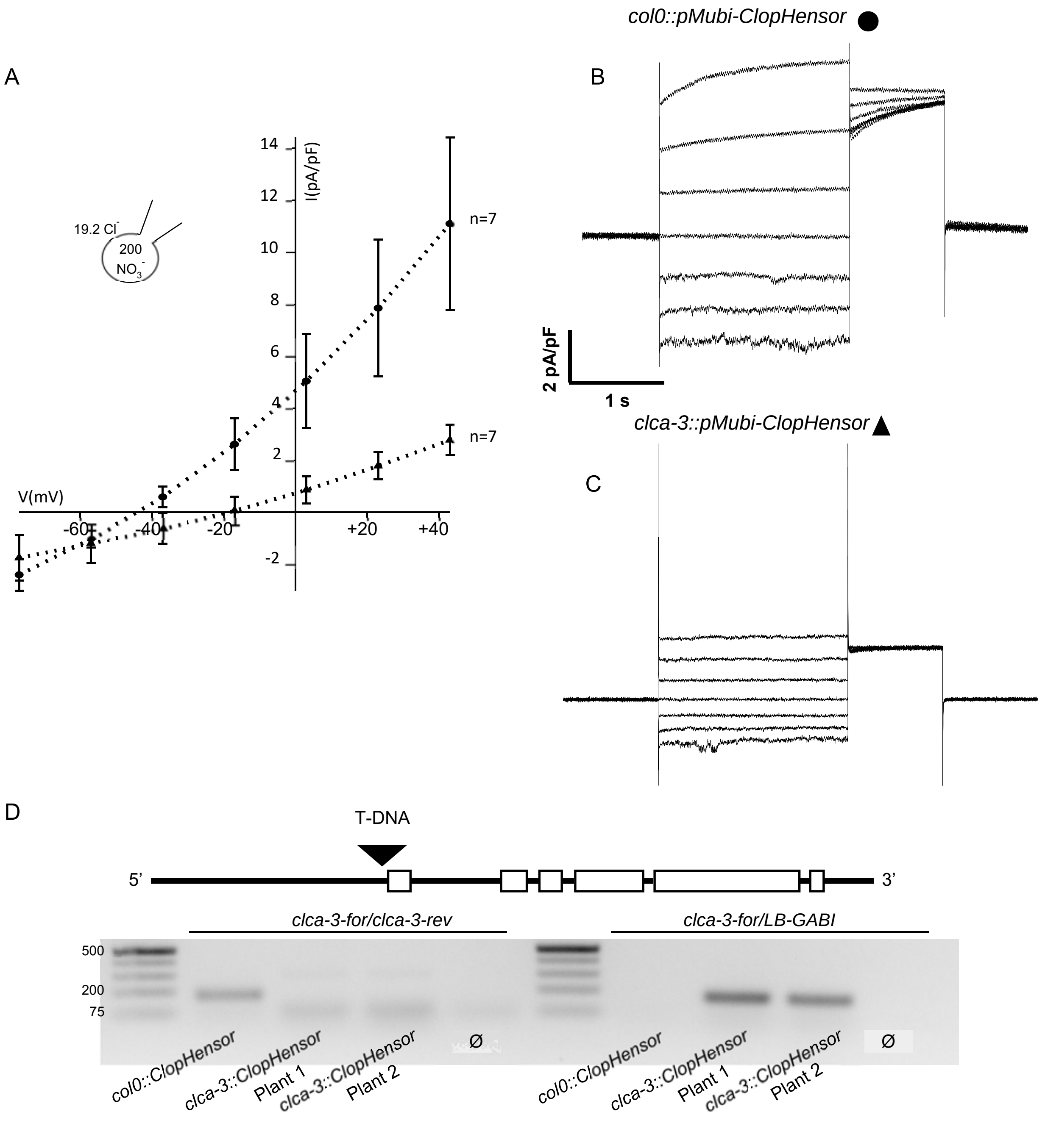


**Fig. S7. AtCLCa mediated vacuolar ionic currents are impaired in *clca-3 pUBI10:ClopHensor* vacuoles.**

(A) Current densities measured in isolated mesophysll vacuoles from wild type and *clca-3* KO plants obtained as described previously (5). Current-voltage curves from patch-clamp measurements in whole-vacuole configuration obtained in vacuoles from wild type *pUBI10:ClopHensor* (circles; n=7) and *clca-3 pUBI10:ClopHensor* mesophyll protoplasts (triangles; n=7). Cytosolic side solution is 7.5mM BTP; 15mM HCl; 0.1mM $CaCl_2$; 2mM $MgCl_2$; 15mM MES at pH 7, osmolarity adjusted to 640mOsm. Vacuolar side solution, 100mM BTP; 200mM $HNO_3$; 1mM $CaCl_2$; 5mM $MgCl_2$; 5mM MES at pH 5.5, osmolarity adjusted at 590mOsm. (B, C) Current traces recorded in wild type *pUBI10:ClopHensor* vacuole (b) and in *clca-3 pUBI10:ClopHensor* vacuole (C). (D) PCR genotypic analysis of *clca-3 pUBI10:ClopHensor* and wild type *pUBI10:ClopHensor.* The primer pairs used were designed to detect wild type allele (clca-3-For/clca-3-Rev) and T-DNA (clca-3-For/LB-GABI). DNA electrophoresis shows that *clca-3 pUBI10:ClopHensor* is homozygous for the T-DNA insertion.

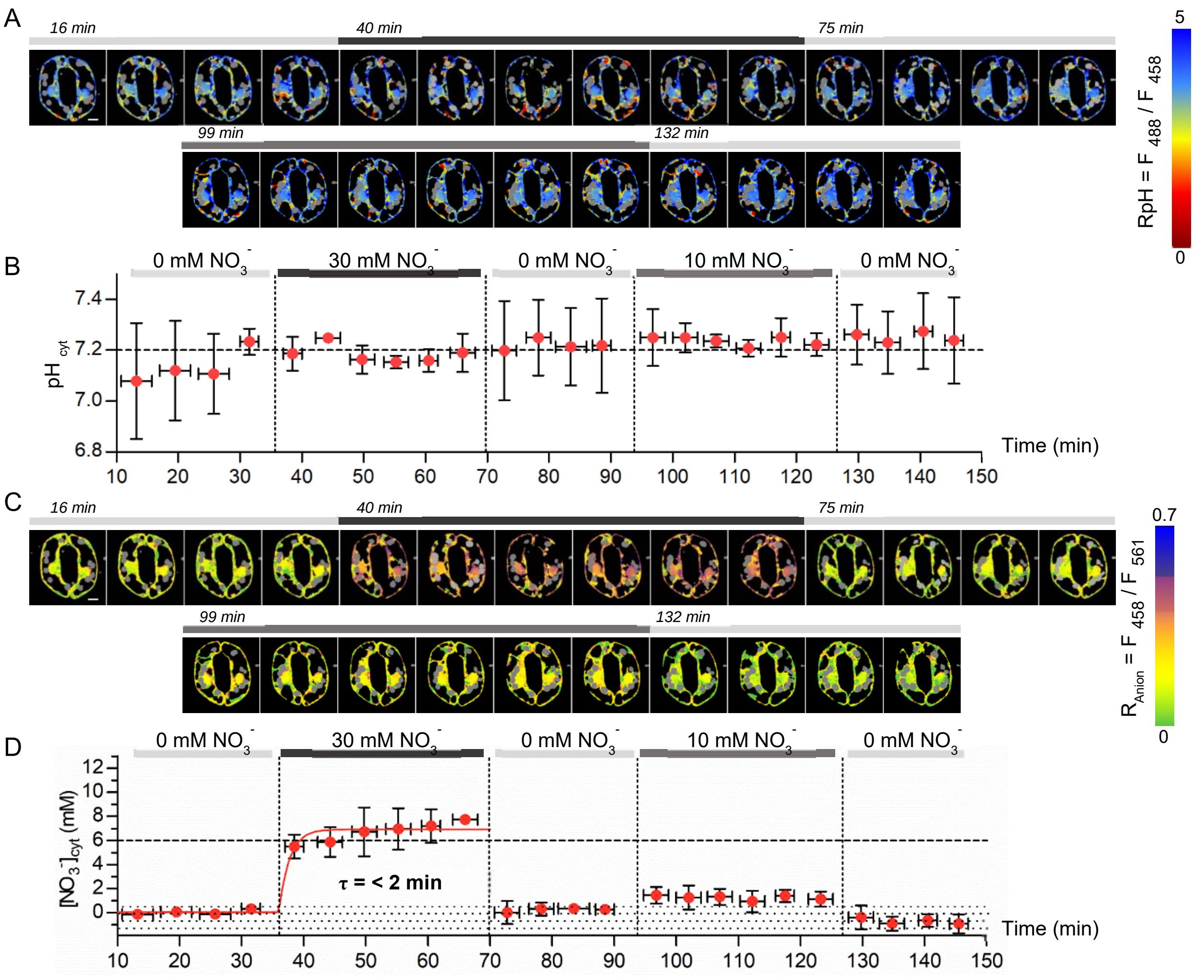


**Fig. S8. The vacuolar $NO_3^-/H^+$ exchanger AtCLCa controls $[NO_3^-]_{cyt}$ kinetics in Arabidopsis GCs.** (A, C) representative false colour ratio images captured every 4 minutes of RpH (A) and $R_{Anion}$ (C) of a typical *clca-3 pUBI10:ClopHensor* stomata. Stomata were exposed sequentially to 30 and 10 mM $KNO_{3,extracell}$. False colours illustrate the different values of RpH (A) and $R_{Anion}$ (C). (B, D) quantification of the mean $pH_{cyt}$ and $[NO_3^-]_{cyt}$ in *clca-3 pUBI10:ClopHensor* stomata from time series like in (A) and (C). (B), mean $pH_{cyt}$. (D), shows the mean $[NO_3^-]_{cyt}$. The line in (D) (red) best fit with equation S6. The $pH_{cyt}$ in each stomata was calculated from equation 1. The $[NO_3^-]_{cyt}$ was calculated with equation 3. The dotted area illustrates the sensitivity threshold for nitrate of ClopHensor. In (B) and (D) each time point is the mean of 3 stomata. Data are mean ± standard deviation. The horizontal bars are the time interval in which the stomata were sequentially imaged. Vertical dotted lines show the time at which the different conditions were applied. Scale bar = 5 µm.

**Supplementary Table 1:**

**Nitrate and chloride content in plantlets**

Colorimetric assay of nitrate and chloride content in Arabidopsis plantlets at different days after germination (DAG); 5 DAG (nitrate n=3; chloride n=1); 8 DAG (nitrate n=3; chloride n=2); 14 DAG (n=5). Each colorimetric assay was performed from 20 to 60 plantlets.

| | **5 DAG** | **8 DAG** | **14 DAG** |
|---|---|---|---|
| **$[NO_3^-]$, mM** | 2.7±0.30<br>(n=3) | 0.96±0.37<br>(n=3) | 0.59±0.11<br>(n=5) |
| **$[Cl^-]$, mM** | 0.76 | 0.035<br>0.058 | - |

### Table S2: Primers list

| | |
|---|---|
| ClopH-For | NNN-GGTACCATGGCTAGCTGGAGCCACCCG |
| ClopH-Rev | NNN-CTCGAGCTACTGGGAGCCGGAGTGGCGG |
| clca-3-For | CACCCACTGTGACTCCTCCT |
| clca-3-Rev | TTTCTCTGCGAATTTTGTCG |
| LB-GABI | TATTGACCATCATACTCATTGC |
